# The low abundance of antimicrobial resistance genes (ARGs) in bacteriophages and their transfer bottlenecks limit the ability of phages to contribute to the spread of ARGs

**DOI:** 10.1101/2024.04.02.587543

**Authors:** Praveen Kant, Bent Petersen, Thomas Sicheritz-Pontén, Kiran Kondabagil

**Affiliations:** Department of Biosciences and Bioengineering, Indian Institute of Technology Bombay, Powai, Mumbai 400076, Maharashtra, India; Center for Evolutionary Genomics, The Globe Institute, Faculty of Health and Medical Sciences, University of Copenhagen, Copenhagen, Denmark; Centre of Excellence for Omics-Driven Computational Biodiscovery (COMBio), Faculty of Applied Sciences, AIMST University, Kedah, Malaysia

## Abstract

The role of bacteriophages in the spread of antimicrobial resistance genes (ARGs) has been debated over the past decade. Several questions regarding the ARG dissemination potential of bacteriophages remain unanswered. For example, what is the frequency of acquisition of ARGs in phages? Are phages selective in acquiring the ARGs compared to other host genes? What is the predominant mechanism of transferring ARGs to phages? To address these questions, we thoroughly analyzed the available phage genomes, viromes, temperate phage, and prophage sequences for the presence of all known ARGs. Out of the 38,861 phage genome sequences we analyzed, only 182 phages contained a total of 314 ARGs. Interestingly, a few of the *Streptococcus* and *Acinetobacter* phages were found to carry an ARG cluster with four or more genes. One of the uncharacterized *Myoviridae* phages was found to carry the entire vancomycin operon. Furthermore, based on the presence of lysogenic marker sequences, the terminal location of ARGs on phage genomes, and complete ARG clusters transferred to phages, we suggest that ARGs are predominantly acquired from hosts by temperate phages via specialized transduction. The close association of most phage ARGs with lysogenic markers and mobile genetic elements (MGEs) also points towards specialized transduction as a potent mechanism of acquisition of ARGs by phages. Our study further suggests that the acquisition of ARGs by phages occurs by chance rather than through a selective process. Taken together, the limited presence of ARGs in phages, alongside various transfer bottlenecks, significantly restricts the role of phages in the dissemination of ARGs.

## Introduction

Genomic and metagenomic studies have shown the pervasive presence of putative antibiotic-resistance genes in the environment [1–4]. Antibiotic resistance can develop either by *de novo* mutations or by the acquisition of antibiotic resistance genes by bacteria from the environment via horizontal gene transfer (HGT). The three well-known mechanisms of HGT are transformation, conjugation, and transduction [5]. Transformation involves the acquisition of naked DNA by a bacterium from its environment, while conjugation entails the direct transfer of DNA from one bacterium to another via the conjugative pilus. The third mechanism, transduction, involves the bacterial virus (bacteriophage or phage)-mediated transfer of DNA from one bacterium to another. Antimicrobial resistance genes (ARGs) are one of the classes of genes that are exchanged between bacteria. Because of the selection of bacteria containing ARGs imparted by the presence of small quantities of antimicrobial residues in the environment, the spread of ARGs via HGT appears to be more rampant than other genes [6]. In the context of dissemination of antimicrobial resistance in the environment, it is important to distinguish between the core bacterial genes that may confer intrinsic resistance (for example, efflux pumps) to the organism under certain conditions and specialty genes that confer direct resistance in the presence of the antibiotic selective pressures (for example, β-lactamases) [7, 8]. Because this distinction is often not made in the literature while annotating genes, especially from the metagenomic studies [9], the reservoir of antibiotic resistance genes (resistome) appears to be much larger than it is. In addition, it is also suggested that the presence of ARGs might have been overestimated due to the less stringent criteria used for their selection [10]. Furthermore, it is hypothesized that several transfer bottlenecks such as ecological connectivity of donor and recipient populations, fitness costs, and pre-existence of resistance in bacterial species in an environment further limit the spread of ARGs in a microbial community [11].

Because of their intricate interaction with their host bacteria, bacteriophages have also been implicated as a major contributor to the spread of ARGs in the environment [9, 12, 13]. Phages mediate DNA transfer between bacteria by either generalized or specialized transduction. In generalized transduction, both lytic and temperate phages aberrantly package host DNA into their capsids instead of their DNA during assembly. Specialized transduction involves only temperate phages which integrate their DNA with the host bacterial DNA. The prophage DNA, due to errors in excision, can pick up a small portion of the host DNA from around its site of excision [14, 15]. Generalized transduction is traditionally considered the primary mechanism of transduction [16–18] with transduction frequencies ranging from 10^-4^ to 10^-7^ [19, 20], while specialized transduction is estimated to be in the range 10^-6^ to 10^-9^ [21, 22]. A higher abundance of ARGs in prophages when compared to free viruses has also been proposed [23]. Though the frequency of lysogeny varies, several reports agree that lysogeny is pervasive in the environment ^2^[24, 25].

Although phages have been shown to mobilize and transfer ARGs [1, 26, 27], they have not been studied comprehensively to gain insights into their potential in ARG dissemination in the environment and the possible mechanisms involved. Here, we report the findings from our comprehensive analysis of phage genomes, viromes, temperate phage, and prophage sequences for the presence of ARGs. Using stringent criteria, we detected the presence of 314 ARGs in 182 phages out of the 38,861 phage genomes. We show that the majority of the ARGs are acquired by phages from their bacterial host via specialized transduction. By correlating the abundance of ARGs in hosts with their phages and the frequencies of horizontal transfer of genes other than ARGs, we suggest that the transfer of ARGs to phages is a non-selective process. We also identify various barriers to the transfer of ARGs through phages. Taken together, our results suggest that bacteriophages play a limited role in the dissemination of ARGs in the environment.

## Results

### Presence of ARGs in phage genomes

As a first step towards understanding the role of phages in ARG dissemination, we identified all ARGs in the phage proteome. A total of 38,861 phage genome sequences from NCBI Virus [28] and GenBank [29] were analyzed. We used the Comprehensive Antibiotic Resistance Database [30] (CARD, a curated and regularly updated database) for identifying the ARGs in phage genomes. In this study, we have included both core bacterial genes that may confer intrinsic resistance to the host and specialty genes that confer direct resistance in the presence of antibiotic selective pressures [2]. Because the mutations that may arise in the core genes as a result of antibiotic selective pressure may also contribute to resistance, the transfer of such genes as well as specialty genes were considered in this study. While the identification of ARGs in phages has been reported widely [1–3, 9, 12, 13], a study suggested that the presence of ARGs might have been overestimated due to the less stringent criteria used for their selection [10]. Hence, we performed a nucleotide-based search (BLASTN) between the CARD database and the phage genome sequences keeping both coverage and identity >= 80%, to rule out any possible false hits and identified the presence of 314 ARGs (Figures 1A). All the hits considered in this study were of high quality with bitscore >=300. Most of the ARGs were related to aminoglycoside (51), macrolide (20), macrolide, and streptogramin (19) classes of antibiotics (Figure 1B). Furthermore, 22 of the ARGs provide resistance against multiple drug classes (ARGs providing resistance against six or more drug classes were considered multiple drug resistant (MDR) ARGs) (Figure 1B). The 314 ARGs identified are from 182 bacteriophages (NCBI panel, Figure 1C).

**Figure 1.**
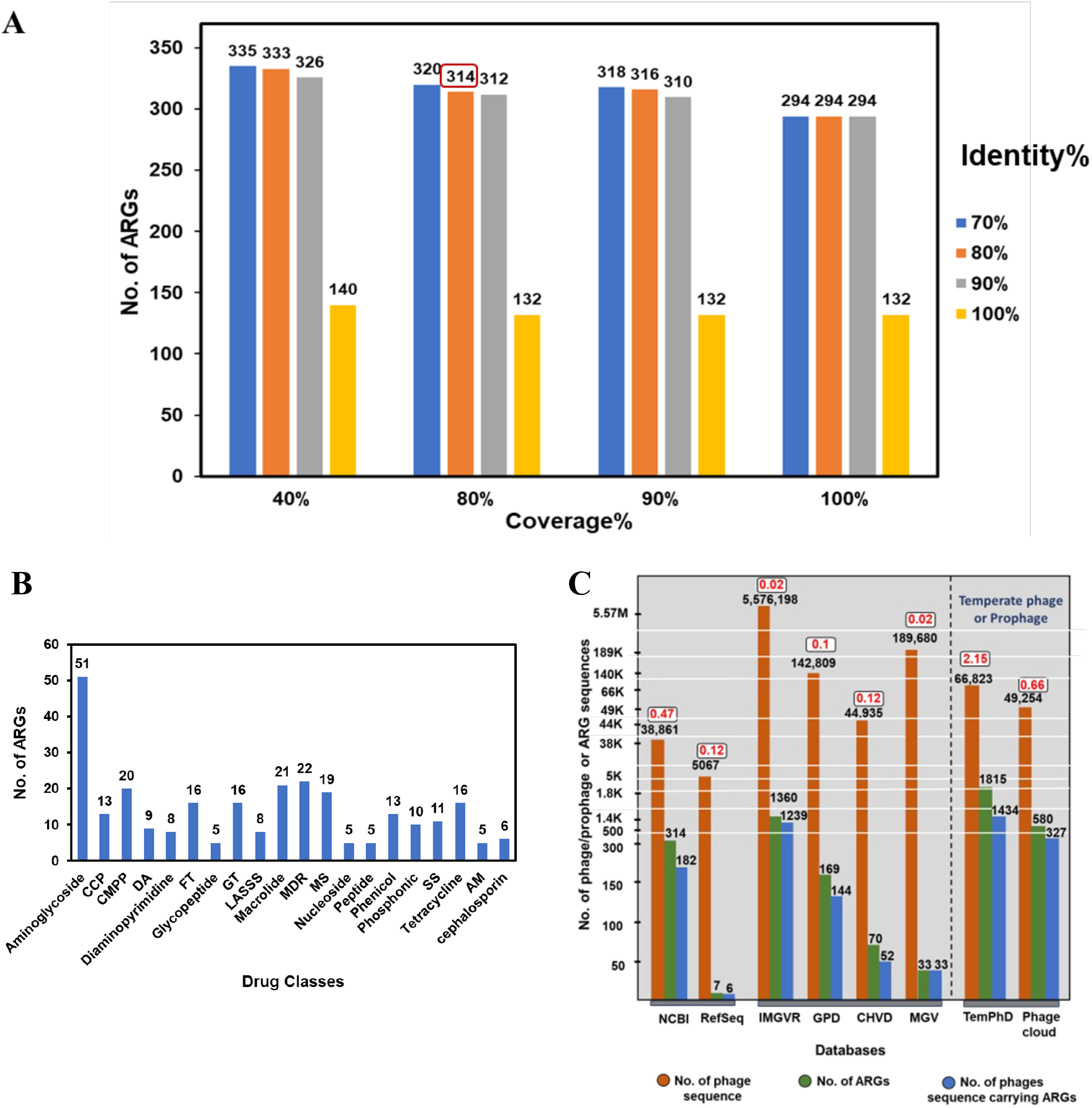
Occurrence of antimicrobial resistance genes/proteins in phage genomes. **(A)** ARGs were predicted by mapping ARGs protein sequence from the CARD (v3.2.8), against bacteriophage protein sequences using BLASTP. The hits generated were filtered at different cut-offs of identity (represented by different colours) and coverage percentages (x-axis). All the hits considered here have a bitscore >= 300. **(B)** Number of ARGs hits for different classes of antibiotics. Antibiotic classes with 5 or more ARGs are shown. CCP: carbapenem; cephalosporin; penam, CMPP: cephalosporin; monobactam; penam; penem, DA: disinfecting agents and antiseptics, FT: fluoroquinolone; tetracycline, GT: glycylcycline; tetracycline, LASSS: lincosamide; macrolide; streptogramin A; streptogramin B; streptogramin, MS: macrolide; streptogramin, SS: sulfonamide; sulfone, AM; antibiotic efflux; macrolide. **(C)** Occurrence of ARGs in various databases. Curated database-NCBI (NCBI Virus & GenBank), RefSeq, virome databases (IMGVR, GPD, CHVD, MGV), and prophage datasets (TemPhD and PhageCloud). The number inside the box in red represents the percentage of phage sequences carrying ARG out of the total phage sequence in the database.

Hosts of most of the ARG-carrying phages were *Acinetobacter* (41.1%), *Streptococcus* (20.4%), and *Escherichia* (10.8%). While no ARGs were found in the 286 complete genome sequences of the *Acinetobacter* phages, 130 ARGs were found in the partial sequences/metagenomic contigs (Supplementary Table S4). About 57% of the hosts of ARG-carrying phages were Gammaproteobacteria, 26.1% belonged to Bacilli, and 0.6% to Erysipelotrichia class (Figure 2, “arc” representing ARGs in phages). The hosts of about 16.2% of ARG-carrying phages are not known.

**Figure 2.**
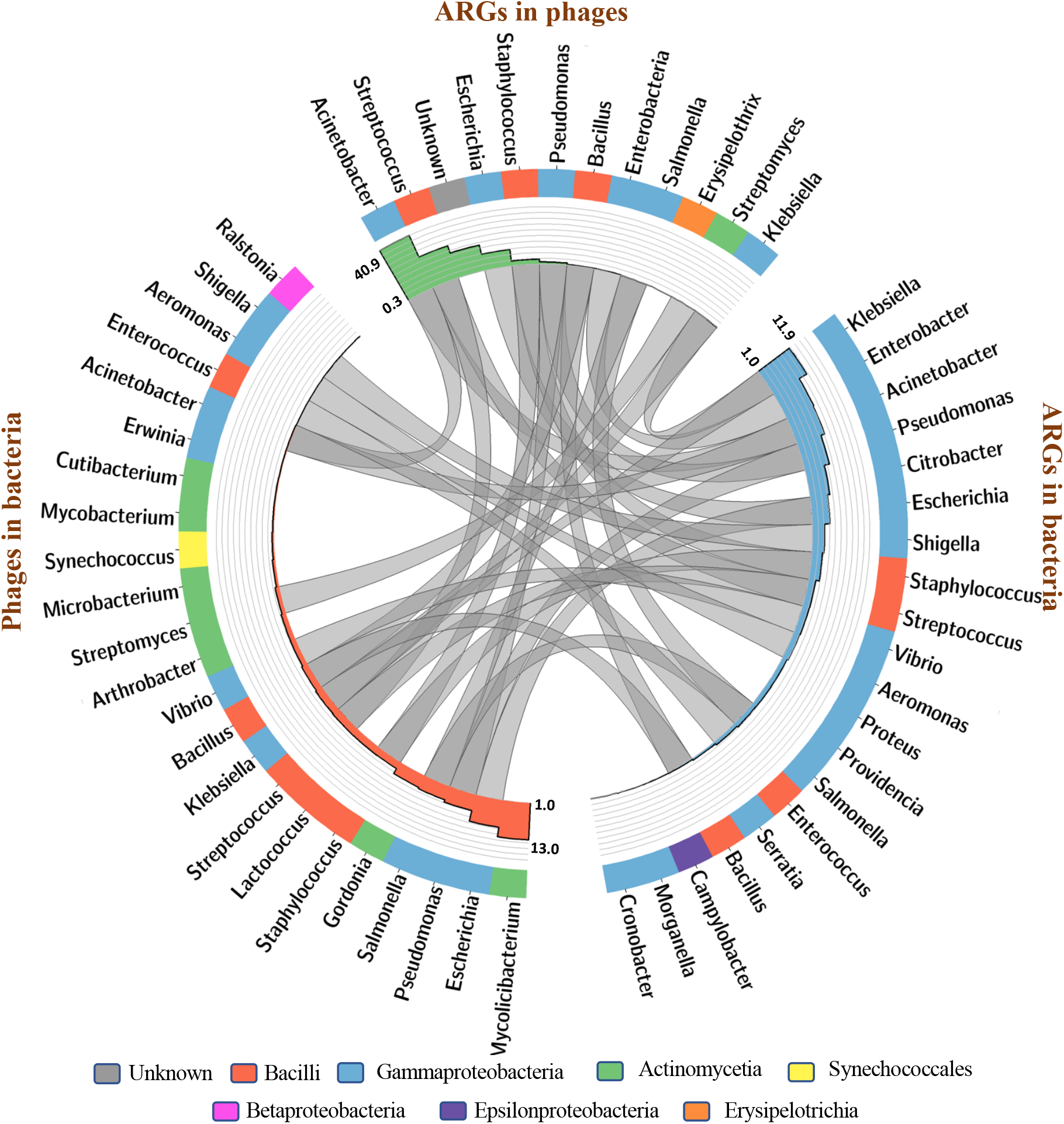
Circos diagram showing the relationships between phages carrying ARGs (ARGs in phages) with ARG abundance in bacteria (ARGs in bacteria) and diversity of phages infecting the bacteria (phages in bacteria). The outer circle is a histogram representing the percentage of ARGs in hosts (green), the percentage of ARGs in different bacterial genera (blue), and the percentage of phages in different bacterial genera (red). Note that each histogram has a different scale. In the core of the circos diagram, the links represent the relationship between ARGs in phages, ARGs in bacteria, and phages in bacteria. Data for phages carrying ARGs was taken from data shown in Figure 1C. Data for ARGs in all bacterial genera was downloaded from CARD Prevalence, Resistome, and Variants data version 3.2.8. A bacterial genus with at least 200 ARGs was considered as ARG abundant. Data for the number of phage species in different bacterial hosts is based on the host information of all the RefSeq bacteriophage nucleotide sequences deposited in the NCBI Virus (https://www.ncbi.nlm.nih.gov/labs/virus/vssi/#/). Bacterial genera with at least 50 phages were considered abundant in phages and included in the study.

Besides NCBI Virus and GenBank, we also searched RefSeq phage genomes (5067) and the following environmental virus databases, IMG/VR [31] (viral (meta)genome repository) (5,576,198), GPD [32] (gut phage database) (142,809), CHVD [33] (viruses from human metagenome associated with chronic diseases) (44,935), MGV [34] (virus from human gut) (189,680), IGVD [35] (infant gut virus database) (10,021), GOV2 [36] (marine virome) (195,699), and STV [37] (soil virome) (4065). In addition, temperate phage and prophage sequences were also screened for ARGs. A total of 66,823 temperate phage sequences were identified using the TemPhD [38] method (fetched from the PhageScope database [39]) and 49,254 prophage sequences extracted from the PhageClouds database [40] (Figure 1C).

Interestingly, a maximum fraction of ARG-carrying phages were found in temperate and prophages sequences, 2.15% and 0.66%, in TemPhD and Phagecloud datasets, respectively, and the least occurrence was found in the MGV environmental database. While only 4 and 1 ARGs were found in the infant’s gut (IGVD) and the marine virome (GOV2) databases, respectively, no ARG was found in the soil virome (STV). In order to estimate the frequency of occurrence of ARGs in bacteriophages with respect to other genes, we used Prodigal to predict the number of genes in phages. We estimated that the frequency of occurrence of ARGs in phages across different datasets ranges from 0 (STV database) to a maximum of 4.5×10^-4^ (temperate phages) (Supplementary Table 1).

### A comparison of the frequency of acquisition of ARGs with respect to other host genes

Our data suggests that *Acinetobacter* (all hits are from the phage partial sequences/metagenomic contigs of *Acinetobacter* phages as mentioned earlier)*, Streptococcus,* and *Escherichia* were the most common hosts of the phages carrying ARGs (arc representing “ARGs in phages” in Figure 2). Furthermore, we compared the transfer frequencies of ARGs with non-ARGs acquired horizontally by phages of these three genera (the dataset used for this analysis was from the “reviewed” and “RefSeq” databases from Uniport and NCBI Virus, respectively, see methods section for details). Our hypothesis here is that if the acquisition of ARGs by phages is beneficial either to the phage or its host, the frequency of transfer of ARGs will be more than other genes. On the contrary, if the acquisition is incidental, the transfer frequencies of ARGs will be comparable to other genes. Out of the total 72098 and 29447 proteins in *Streptococcus* and *Escherichia genera* respectively, 187 (0.26%) and 600 (0.68%) annotated proteins have been transferred to their respective phages; and 63 and 24 are the ARGs transferred between the two host-phage pair. Similarly, only 4 annotated non-ARGs out of the total 52241 genes were found to be exchanged between the *Acinetobacter* genus and its phages. We categorized all the horizontally transferred non-ARGs as Clusters of Orthologous Groups (COGs) (Figure 3). Genes with similar functions are usually adjacent to each other and they have a higher probability of moving together [41, 42]. Most of the genes transferred to phages of the two bacterial genera are uncharacterized. In *Streptococcus*, though ARGs are one of the most abundant groups transferred, their frequency of acquisition is comparable to the COG labelled as L (“replication and repair”). Besides, G (“carbohydrate metabolism and transport”) was the second most abundant COG. Similarly in Escherichia, COGs L, K (transcription), and M (Cell wall membrane/envelope biogenesis) were the most abundant (Figure 3). This result shows that there is no specific bias in the acquisition of ARGs by phages of both *Streptococcus* and *Escherichia* genera. The transfer of more genes from the *Escherichia* genus to their phages, when compared to *Streptococcus*, appears to be simply a function of the genome size and the number of genes (the *Escherichia* genome is more than twice the size of the *Streptococcus*).

**Figure 3.**
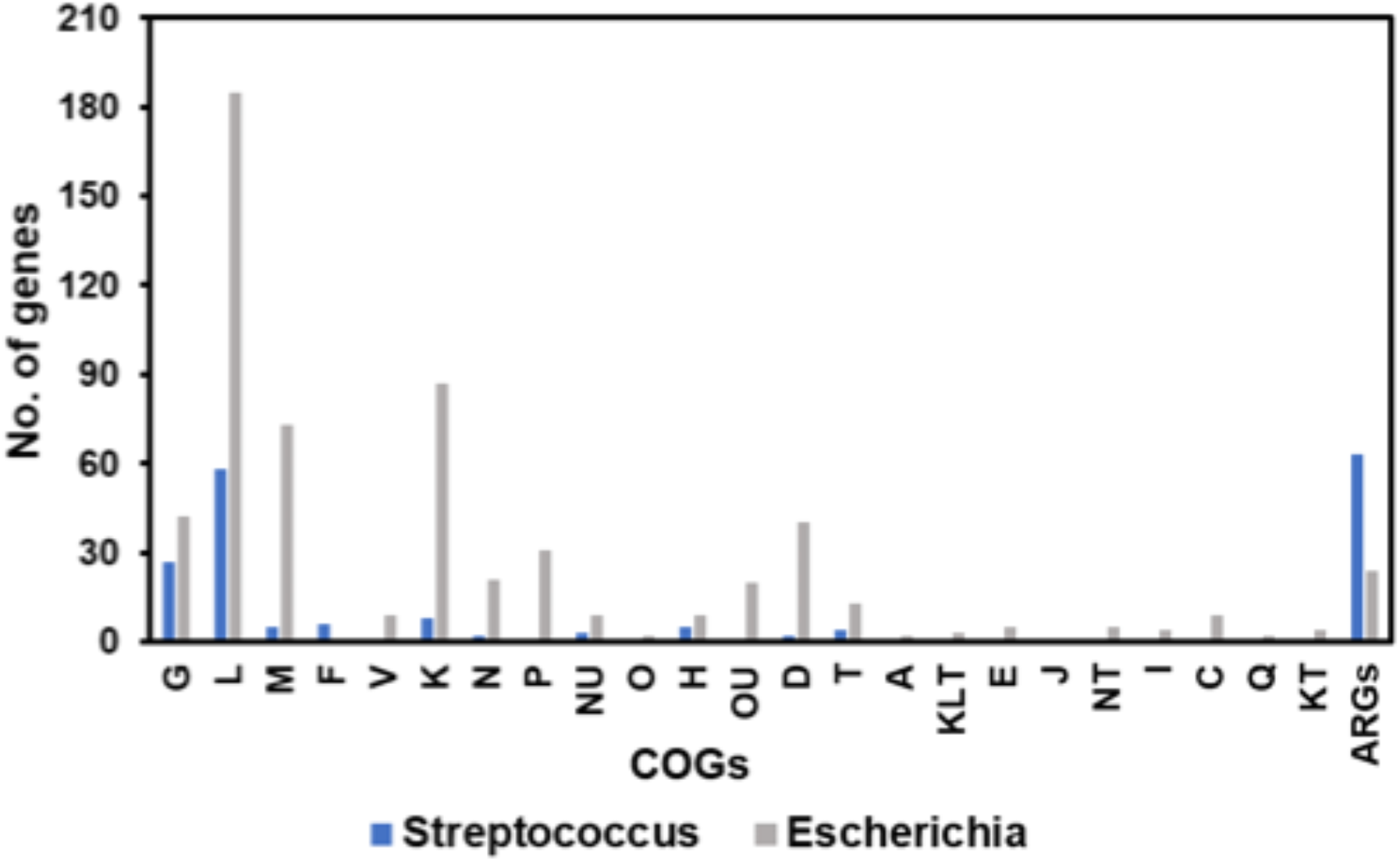
A comparison of the number of horizontally transferred COGS and ARGs from *Streptococcus* (in blue) and *Escherichia* (in grey) with their respective phages. Horizontally transferred genes in the respective phages of two bacteria were identified by mapping the proteome of phages with bacterial proteome (identity > 80%, coverage > 80%, evalue < 10^-5^). These genes were later classified into different COGs (Cluster of Orthologous Groups). In both the bacteria most of the genes transferred to phages are uncharacterized (not shown; 146 and 15057 genes in *Streptococcus* and *Escherichia* respectively). Abbreviations used for COGs -A: RNA processing and modification, C: Energy production and conversion, D: Cell cycle control; cell division and chromosome partitioning, E: Amino Acid metabolism and transport, F: Nucleotide metabolism and transport, G: Carbohydrate metabolism and transport, H: Coenzyme metabolism and transport, I: Lipid metabolism and transport, J: Translation; ribosomal structure and biogenesis, K: Transcription, L: Replication; recombination and repair, M: Cell wall/membrane/envelop biogenesis, N: Cell motility, O: Post-translational modification, protein turnover, chaperone functions, P: Inorganic ion transport and metabolism, Q: Secondary metabolites biosynthesis; transport and catabolism, S: Function Unknown, T: Signal Transduction, U: Intracellular trafficking and secretion, V: Defence mechanisms.

### Correlation between the abundance of ARGs in phages with abundance of ARGs in bacteria and the number of phages of bacteria

Furthermore, we examined if the abundance of ARGs in phages correlates with the abundance of ARGs in their respective bacterial genus, as well as the abundance of phages infecting a particular bacterial genus (Figure 2). If the transfer is random, ARGs would be transferred based on the abundance i.e., the hosts carrying a higher number of ARGs and bacteria with more phages, the respective phages would acquire more ARGs.

Although the median number of ARGs in bacterial genera was 11, to assess the most ARG- abundant bacteria, we considered the top 1% bacterial genera (200) of the total ARGs (20041). Out of the 166 bacterial genera carrying ARGs, 20 were found to meet this criterion (Figure 2, Supplementary Table S2). About 86.3% of the ARGs present in these ARG-abundant bacteria belong to the gammaproteobacterial class, 12.2% belong to the bacilli class, and the rest (about 1.4 %) to epsilonproteobacteria (Figure 2, arc ARGs in phages). From this data, it is clear that the ARGs are mostly carried by a very small group of bacteria. For this study, a bacterial genus with at least 1% (54 phages) of the total phages were considered phage-abundant bacteria. Out of the 187 bacterial genera carrying ARGs, 23 were found to meet this criterion (Figure 2, Supplementary Table S3). Bacterial genera with a maximum number of phages are *Mycolicibacterium, Escherichia*, and *Pseudomonas* in that order (Figure 2, arc—phages in bacteria). Most phage-abundant bacteria belong to class Gammaproteobacteria (43.3%), Actinomycetia (35.9%), Bacillus (18%), and Synechococcales (2%).

Our results suggest that gammaproteobacteria and bacilli are among the bacteria that carry the highest number of ARGs and are also hosts to a larger number of phages. Incidentally, phages of these bacteria carry more ARGs than other phages in the database. This association suggests that the process of transfer of ARGs from host bacteria to phage is unlikely to be selective.

### Role of specialized transduction in ARG acquisition in phage

We also studied the genomes of phages for possible markers of lysogeny. Lysogeny is usually mediated by site-specific recombination systems [43, 44] that employ a diverse set of components. We included all of them in our study, namely, *int* (integrase)/recombinase (serine and tyrosine recombinase) [45], *xis* (excisionase), *cI* (repressor – represses expression of all other phage genes), *cII* (activator – for *cI* and *int*), *cIII* (protects *cII* from degradation), and *attP* (attachment site in phage genome) [44, 46, 47]. Of them, integrase/recombinase appears to be the most common marker of lysogeny. The presence of one or more of the above markers was considered a signature of lysogeny [46, 48, 49]. We find that a majority (60%) of the phages carrying ARGs are temperate phages. If we exclude the partial sequences where lysogeny markers could not be confirmed, the fraction of temperate phages goes up to 92.5 % (Supplementary Table S4). Interestingly, except FN997652.1 (streptococcus phage), 14 out of the 22 phages carrying 4 or more ARGs were found to be temperate phages, and 12 were partial genomes, leading us to hypothesize that these phages might have acquired ARGs via specialized transduction.

To check this, we located the position of the ARGs in the complete phage genomes. We found that 64 ARGs (out of 115) were present towards the genome termini (Figure 4A). In general, the lysogeny markers like integrase, excisionase, and attP are located towards the phage genome terminal and help in phage genome integration and excision during lysogeny [46, 50]. The phage ARGs were found to be co-localized with lysogenic markers, specifically integrase, and distributed similarly in the genome (Figure 4A). In temperate phage and prophages data from two sources TemPhD and PhageCloud 60% and 50.4% of the ARGs were found to be distributed at the terminals (Figure 4B).

**Figure 4.**
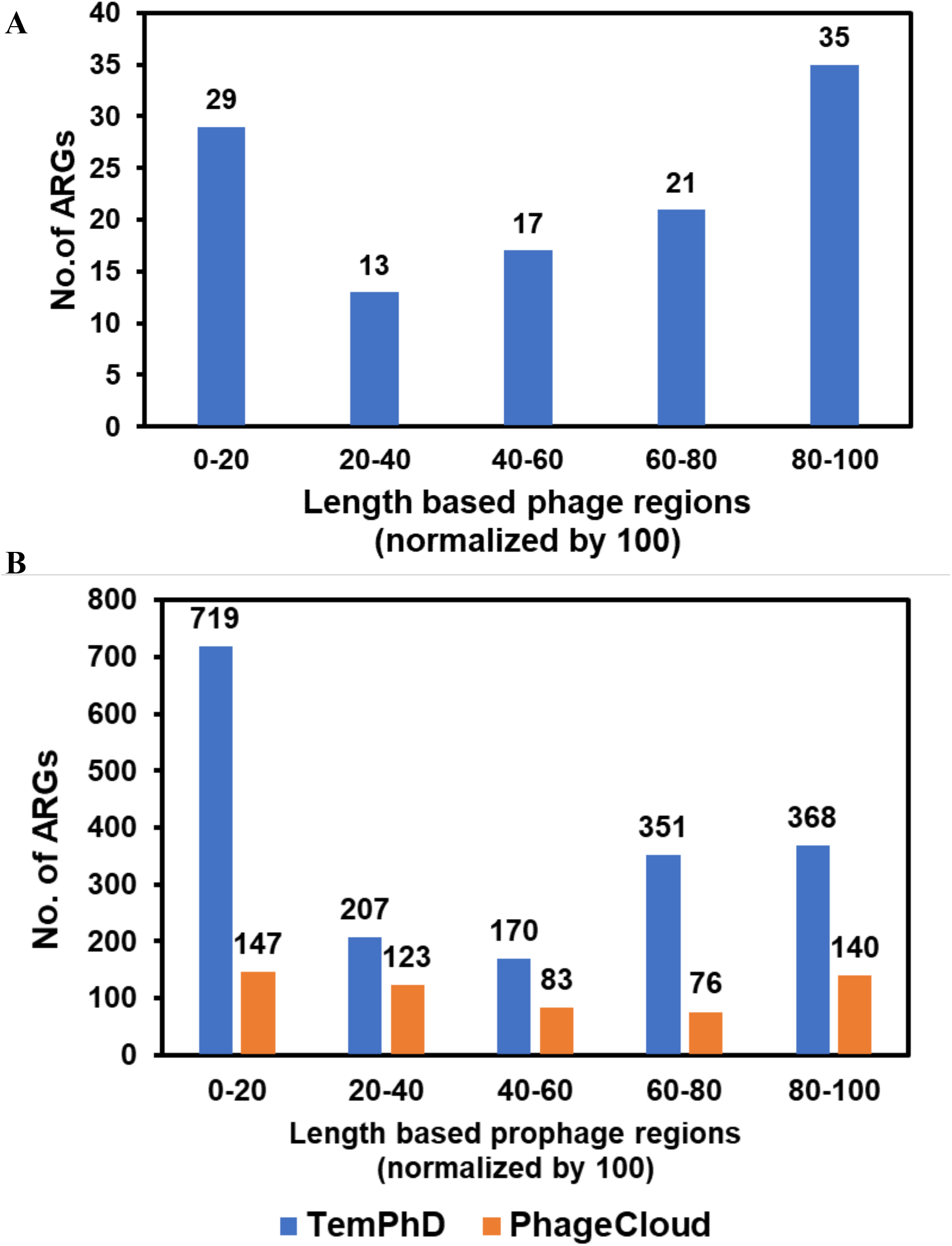
Location of ARGs in phage genomes. **(A)** Genomic distribution of ARGs, Lysogeny marker sequences, and MGEs in the phage genome. (**B)** ARG distribution in prophage genomes.

We investigated the genes in the flanking regions (5 kb upstream and downstream) of the phage ARGs and found colocalization of mobile genetic elements (MGEs) and ARGs in ∼44% of the phages (Supplementary Table S4). MGEs help in the translocation of genes within or between genomes. Phage would have acquired these MGEs along with ARGs. Their presence is considered to increase the dissemination potential and the “risk factor” of ARGs [7]. The MGEs found were Tnp3, tn1270, IS26, IS481, tnpA, ISSod13, IS605, TnpW, IS1216, tn5252, IS903B, TnpR, Mos transposase, IS256, IS257, TnpV (http://124.239.252.254/danmel/index.php) [51].

Touchon et. al. [52] reported that bacteria with fast doubling times carry more prophages. We found that a majority of phages carrying ARGs in our dataset were temperate phages (Supplementary Table S4). The hosts of these temperate phages turned out to be fast-growing pathogenic bacteria when we compared our dataset with the dataset used by Touchon et al. (Supplementary Table S5). This result suggests that temperate phages are selected over non-temperate for the fast growth of the host. Thus, they increase the chances of ARG acquisition from the host due to the increased genomic interaction and exchange between phage-host pairs via specialized transduction.

### Multiple ARGs present in phages are part of the gene cluster

If four or more ARGs were present together in close proximity and together responsible for conferring resistance to single or multiple antibiotics we considered them as an ARG cluster. From our analysis, the close proximity is approximately within the ∼20% of the genome. We identified four clusters, one each in a Streptococcus and Acinetobacter phage (from metagenomic contigs), one each in the Escherichia virus P1, and an unclassified Myoviridae phage (Figure 5). The Streptococcus phage cluster (APH(3’)-IIIa, SAT-4, and (6), ErmB), was found in the 4 streptococcus phages out of the total 878 Streptococcus phages. These clusters are identical and present towards the end of the genomes suggesting that the transfer might have happened only once (Figure 5). The same cluster is also present in the probable hosts, *S. suis, S. dysgalactiae, S. acidominimus* and *S. oriscaviae* (Supplementary Table 6).

**Figure 5.**
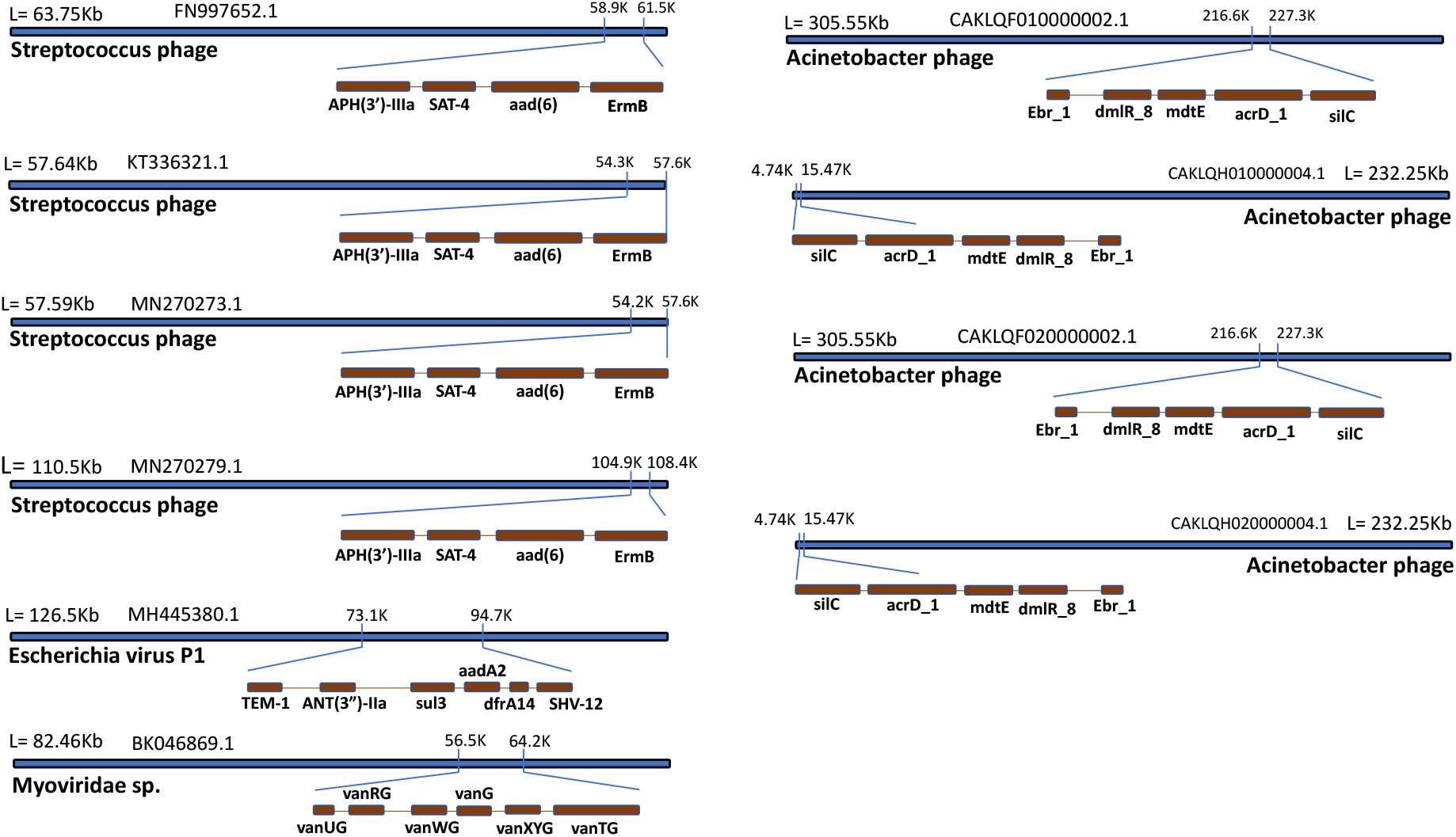
ARG clusters in phage genomes. If 4 ARGs are present together in close proximity (∼20% of the genome) and together responsible for conferring resistance to single or multiple antibiotics we considered them as an ARG cluster. The ARG clusters were found in Streptococcus phages, Escherichia phage, Myoviridae, and Acinetobacter phages.

The cluster present in Escherichia virus P1 is comprised 6 different ARGs providing resistance against 4 drug classes (Figure 5). Although this cluster is present towards the middle of the genome, since all ARGs are present together, it is most likely that the transfer of the cluster has happened via specialized transduction. Similarly, the Myoviridae cluster is also present towards the middle and contains a complete operon of vancomycin resistance [53, 54] (Figure 5). As far as we know, this is the only example of a complete ARG operon acquired by a phage. Further, in *Acinetobacter* phage partial sequences/contigs, two types of 4 ARG clusters were identified, one towards the terminal and the other towards the middle region.

### Source of ARGs

Additionally, we examined the prevalence of all the known ARGs in plasmids and genomes to assess whether a bias exists in the frequency of occurrence of ARGs on genomes as compared to plasmids. We expected a higher prevalence of ARGs on genomes than plasmids since the interaction of phages with plasmids appears to be limited [55] and hence phages would predominantly acquire ARGs from the bacterial genome. About 66 % of the ARGs in the CARD database have the highest prevalence in whole genome sequencing of bacteria, with 25% showing prevalence in genomic origin and the remaining 9 % in plasmid (Figure 6A). Further, only 4 % of all ARGs in the CARD database are mapped with the Plasmid Database (PLSDB, Figure 6B) suggesting that most of the ARGs are present in the bacterial genome. In our study also only 3% of ARGs found in phages have their prevalent origin in a plasmid (Supplementary Table S7). Since viruses interact predominantly with the host genome during lysogeny^54^, the high prevalence of ARGs in the genome compared to plasmids increases the chances of phages acquiring ARGs from the host genome.

**Figure 6.**
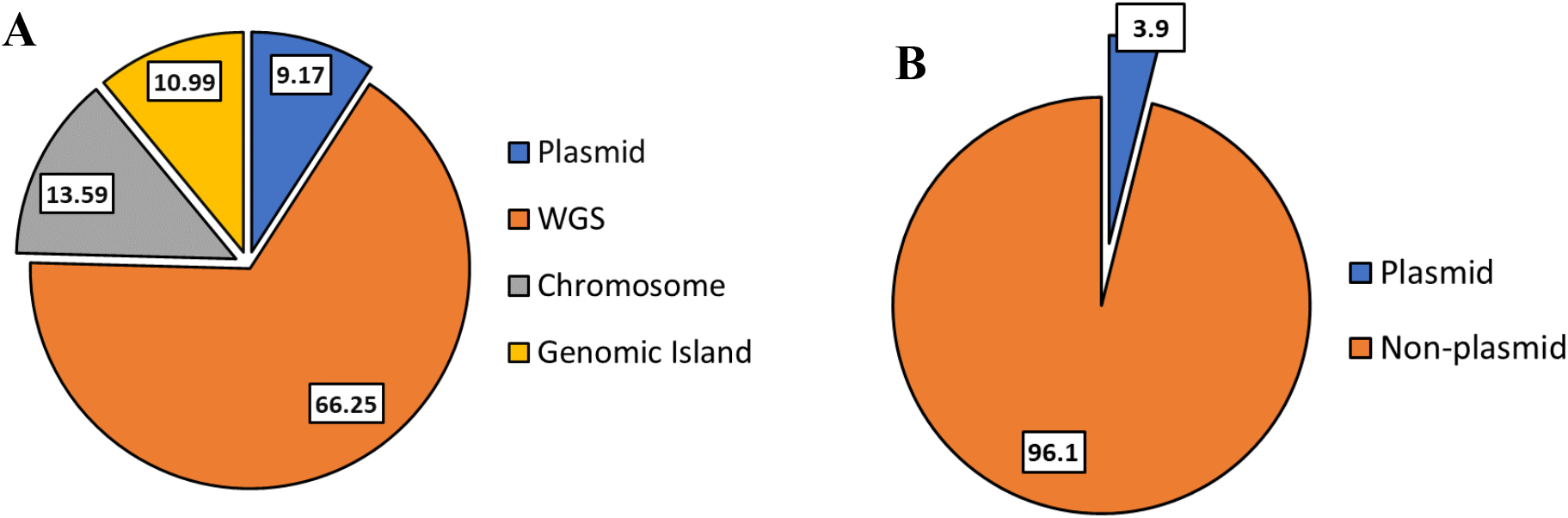
Source of ARGs. **(A)** CARD prevalence data was used to find the probable source of ARGs, whether they are located in the chromosome or plasmid. **(B)** The same dataset was also compared with the Plasmid database (PLSDB).

## Discussion

Bacteriophages, because of their unprecedented abundance and diversity, are thought to play a major role in shaping environmental microbial communities. Studies suggest that bacteriophages may kill approximately 20 % of bacteria in oceans, influence their metabolism, and also mediate horizontal gene transfers between bacteria [56, 57]. In the past decade, the availability of huge amounts of genomic and metagenomic data has enabled studies on phage-mediated horizontal transfer of genes between bacteria, frequencies of their transfer, and the underlying mechanisms [3, 9, 58–61]. An important question researchers are trying to answer is how significant is the role of phages in the spread of ARGs. While some studies downplay their role [62], others suggest that phage-mediated transduction may play a substantial role in the spread of ARGs [3, 9, 12, 13, 63]. In this study, we strive to find cues to this question by carrying out a comprehensive analysis of available phage genomes, RefSeq genomes, prophages, and viromes.

We used sequence alignment-based search yielding high-quality results (query coverage – 80%, and identity – 80%) that show the presence of only 314 ARGs in 182 phages (out of 38,861 phage genomes, Figure 1). Our results further suggest that gammaproteobacteria and bacilli are among the bacteria that carry the highest number of ARGs and are also hosts to a larger number of phages. While a direct correlation between antibiotic exposure and the abundance of ARGs in viromes has been suggested previously, the same is not known for viral genomes. Incidentally, there is a strong association between the ARG abundance in phages with the ARG abundance in host bacteria and the number of phages infecting the bacteria (Figure 2). We identified and compared bacterial genes other than ARGs that are transferred horizontally in *Escherichia,* and *Streptococcus* genera, as these two were among the most abundant in ARGs and also hosts to most phages carrying ARGs (Figure 3) and found no specific bias in the acquisition of ARGs by phages of both genus. The transfer of more genes from bacteria belonging to the *Escherichia* genus to their phages, when compared to *Streptococcus*, appears to be simply a function of the genome size and the number of genes (the *Escherichia* genome is more than twice the size of the *Streptococcus*).

We also analyzed the genomic context of the ARGs present in phages and found that a majority of phages (92.5 %, excluding partial sequences where lysogenic markers could not be confirmed, Supplementary Table S4) are associated with lysogenic markers and most of the ARGs are present towards the genome termini suggesting specialized transduction as the potent mechanism of transfer of ARGs (Figures 4). Consistent with this observation, we found that temperate phages and prophages harbor about 6 and 1.9 times more ARGs respectively when compared to all phage sequences (from NCBI Virus and GenBank) (Figure 1C: TemPhD and PhageCloud with respect to NCBI panel). Further, only of small number of ARGs among all the CARD ARGs have their origin from plasmid (Figure 6A), and also among the ARGs found in phages, only 3% have prevalence in plasmid (Supplementary Table S6). A study by Meng et al also suggested that most ARGs are present in the bacterial genomes rather than plasmids [64]. While plasmid-mediated transfer of ARGs between bacteria, especially in soil, has been implicated in ARG dissemination [65], the chances of phages acquiring ARGs (or other genes) from plasmids appear to be rather limited. Although it is reported that the presence of plasmids influences phage infectivity [66], it is not clear whether the phage genomes interact with plasmids and what would be the outcome of such interactions. Based on our observations from this study, we hypothesize that phage genomes rarely interact with plasmids during their replication and assembly cycle.

Traditionally, generalized transduction has been suggested as the main mechanism of acquisition of ARG by phages [3, 10, 15, 59]. While we can’t rule out the role of generalized transduction in the transfer of ARGs between different bacteria, from this study, the presence of lysogeny markers in phages carrying ARGs, terminal abundance pattern, and the colocalization of ARGs and MGEs, high abundance of ARGs in host chromosome with respect to plasmid; and the transfer of complete ARG clusters to phages suggests that the specialized transduction is a prevalent mechanism of acquisition of ARGs (or any other genes) by phages. Site-specific recombination is the core mechanism in specialized transduction for the interaction between phage and bacteria, other recombination mechanisms like intergenic – between two different phages and between host-phage (P2), intragenic (in phage T4 [67] and transposition in phage Mu [68]) can occur but their contribution to overall gene acquisition by phages could not be assessed in this study.

While generalized transduction frequencies are reported to be in the range of 10^-4^ to 10^-7^ [19, 20], specialized transduction is estimated to be in the range of 10^-6^ to 10^-9^ [21, 22]. Both modes of gene transfer have several limitations. In generalized transduction, due to the presence of pseudo-pac sites, host genome fragments are accidentally packaged [14]. In most cases, these fragments lack phage genes for defence, replication, and recombination, and hence are degraded by the host immune system [69] limiting the gene transfer via generalized transduction to similar species/strains. Specialized transduction is limited to the transfer of host genes present near the prophage genomes. Transfer of genes by both modes of transduction to the bacterial community is also limited by the host range of the phages.

It is also reported that the frequency of lateral transduction, which is a type of specialized transduction [14, 60], is at least 1000-fold more than the normal “excision-replication-packaging” mode of specialized transduction. In the case of lateral transduction, prophage undergoes theta mode of replication and delayed excision resulting in high-frequency in-situ packaging of prophage transforming the bacterial region into a hypermobile region of gene transfer [56]. Considering the intrinsic limitations of gene transfer by generalized transduction and the recent discovery of the lateral transduction phenomenon, specialized transduction appears to be a potent mechanism for phage-based gene transfer.

Genes are generally clustered based on their involvement in the same metabolic pathway, genes encoding interacting proteins or genes that affect the same trait [70] and it is a common feature of bacteria [71]. Clustering also helps in the co-transfer of genes horizontally passing on the complete functions during evolution [42, 72]. Examples include PhD/Doc system of bacteriophage, restriction/modification systems, and ARGs [41]. In the case of ARGs, co-clustering provides the host bacteria resistance against diverse antibiotics. We found the presence of 10 ARG clusters of 5 different types and interestingly, in one case, the complete operon of vancomycin. Furthermore, 4 ARG clusters were found to be associated with MGEs. Since all ARGs are present together, it is most likely that the transfer of the cluster has happened via specialized transduction (Figure 7A).

**Figure 7.**
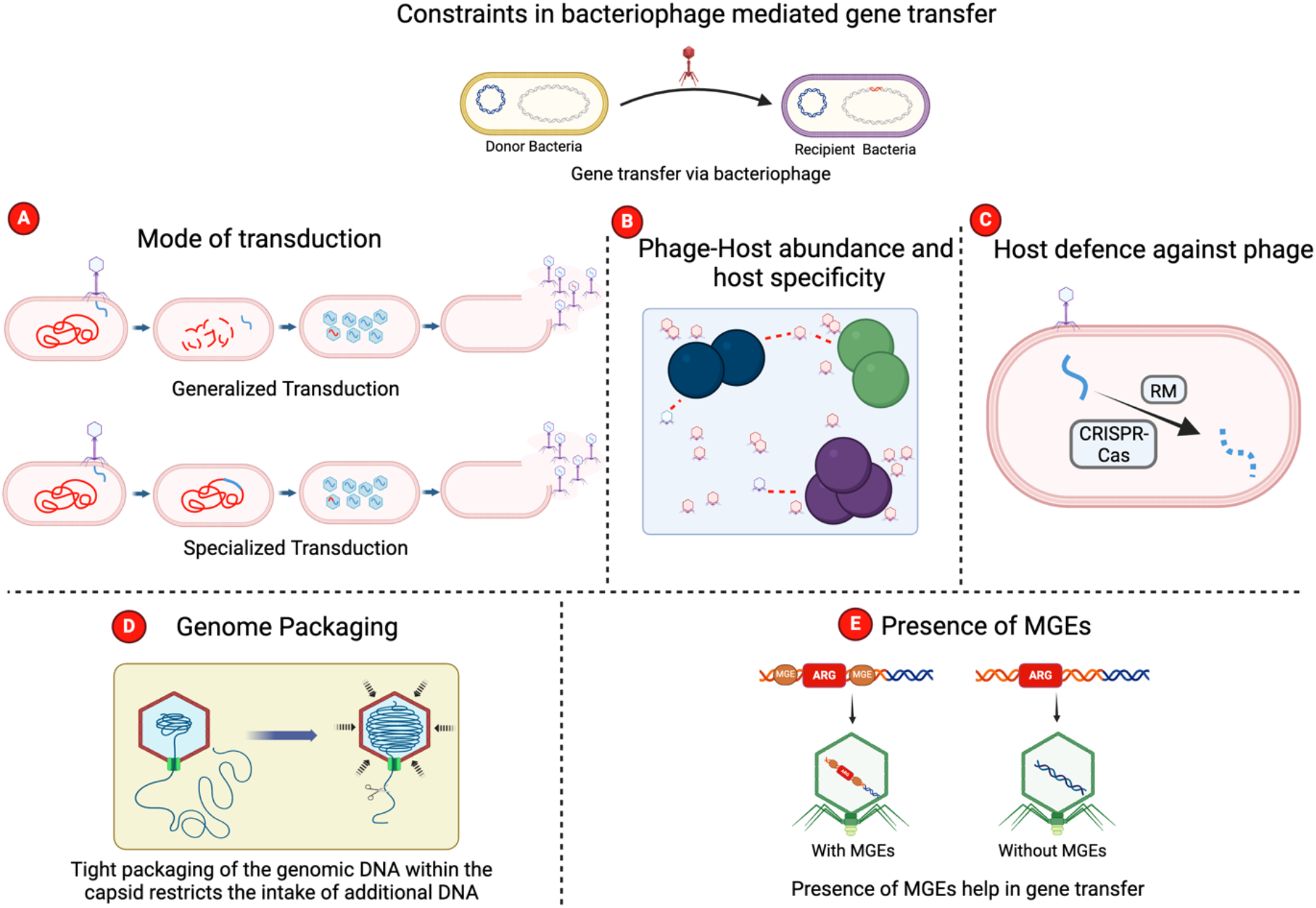
Schematic representation of constraints in bacteriophage mediated gene transfer. (A) Mode of transduction. Our study suggests specialized transduction as a predominant mechanism of gene acquisition by bacteriophages. (B) Phage and host abundance and their spatiotemporal dynamics influence the number of interactions between phage and bacteria. Further, the phage’s host range also limits the phage-host interactions. Condition-dependent lysogeny may also influence the lateral transfer of host genes. (C) Host defence systems prevent the adsorption of phage and entry of phage’s DNA into the host. After entry, systems such as CRISPR-Cas and restriction-modification system (RM) limit the phage’s success. (D) Tight packaging density and limited phage capsid volume limit the acquisition of extra DNA. (E) In the absence of Mobile Genetic Elements (MGEs), the transfer of flanking genes/ARGs from host to phage genome and its dissemination to other bacteria is constrained. The illustration was created using the BioRender software.

Since the estimated number of phages in the biosphere 10^30-31^ [12] is much more than the phages analyzed in this study, the actual number of ARGs and other bacterial genes transferred would be conceivably significantly more than what our results suggest; however, we believe that the frequency of transfer would be similar to what our results suggest. It is also estimated that viruses in the oceans produce about 10^23^ infections every second [73]. While it is possible that such extensive interaction with the microbial communities in many ecosystems, in addition to microbial mortality, will have many plausible outcomes, including horizontal transfer of genes between and across phages and their hosts, phylogenetic barriers [74] in such niches severely limit the exchange of genes, especially ARGs. For phage-based gene transfer to occur the first step is that the two, bacteria and phage, should interact with each other, which depends on their abundance. While phages are estimated to outnumber bacteria by at least 10-fold [73, 75], the host specificity and the very low probability of temperate phages acquiring ARGs/ARG clusters in an aquatic niche further limits their spread via phages (Figure 7B). In addition, a large number of host defence mechanisms (receptor binding proteins, Resistance-Modification systems, CRISPR-Cas, etc.) also significantly limit the success of phages (Figure 7C). Another factor limiting the transfer of ARGs from bacteria to phages and back is the limited capacity of the temperate phages to accommodate additional DNA. While some temperate phages can acquire up to ∼ 5% additional DNA [19], such acquisitions are rare as shown in this study (Figure 7D). Further, dissemination of ARGs will also depends on the presence of MGEs (Figure 7E). Taken together, our result suggests that bacteriophages play a limited role in the dissemination of ARGs in the environment.

## Material and Methods

### Data collection and ARG identification

CARD (4.0.2-Nov 2023) [30] with 4804 ARGs was used for this study. Bacteriophage nucleotide sequences were fetched from the NCBI Virus^41^ and NCBI GenBank (Nov 2023). 38,861 unique phage sequences were fetched from these two repositories. Following phages sequences from different environments were also considered: IMG/VR [31] (5,576,198), GPD (142,809) [32], CHVD [33] (44,935), MGV [34] (189,680), IGVD [35] (10,021), GOV2 [36] (195,699), and STV [37] (4065). In addition, temperate phage and prophage sequences were also screened for ARGs. A total of 66,823 temperate phage sequences were identified using the TemPhD [38] method fetched from the PhageScope database [39] and 49,254 prophage sequences extracted from the PhageClouds database [40].

Traditionally, for identifying ARGs, a threshold of 25 amino acid coverage and 90% identity were considered [76, 77]. These thresholds were too lenient resulting in false positives and over-estimation of ARGs [10]. Hence, at least 40% coverage with 80% identity of nucleotide is proposed as the conserved threshold for ARG prediction [78]. We performed standalone BLASTN [79] for ARG prediction in bacteriophage and employed a stringent criteria of threshold of 80% for both coverage and identity and e-value 10^-3^.

### HGT and the correlation study

We compared the transfer frequency of other bacterial genes (referred to as non-ARGs in this study) with ARGs. *Acinetobacter*, *Escherichia,* and *Streptococcus* were used for this study as most of the phages carrying ARGs infect these three bacterial genera. Proteomes of the three bacterial genera were fetched from the UniprotKB Proteome database [80]and their respective phage RefSeq proteins were obtained from the NCBI Virus database (https://www.ncbi.nlm.nih.gov/labs/virus/vssi/<u>#/</u>) [28]. We performed BLASTP between bacterial proteome and their respective phage proteins with 80% as the cutoff for both identity% and coverage%. Characterization of these sequences was done using EggNOG-mapper [81].

To check the correlation between the abundance of ARGs in phages with the abundance of ARGs in their respective host bacterial genus, as well as phages infecting a particular bacterial genus, data for ARG abundance in all bacterial genera were downloaded from the “CARD prevalence, resistome and variants” database version 4.0.2. To know the number of phage species in a different bacterial host, we downloaded host information of all RefSeq bacteriophage nucleotide sequences deposited in the NCBI Virus database (https://www.ncbi.nlm.nih.gov/labs/virus/vssi/<u>#/</u>).

We kept a high cut-off for both parameters. Bacteria with 200 (1%) ARGs were considered ARG abundant and bacteria with 50 (1%) and above phage were considered abundant in phages. Relationship between the three parameters: the abundance of ARGs in phages, abundance of ARGs in bacteria, and number of phages infecting bacteria, were represented using the Circos software [82].

### Identification of the mechanism of transfer

We used the following gene/DNA segments as markers for lysogeny: *int* (integrase)/recombinase (serine and tyrosine recombinase, *xis* (excisionase), *cI* (repressor – represses expression of all other phage genes), *cII* (activator – for *cI* and *int*), *cIII* (protects *cII* from degradation) and *attP* (attachment site in phage genome). We identified lysogeny markers manually searching the phage genomes carrying ARGs and also performed standalone BLASTN [79] between the lysogeny markers as query against the phage sequences with the following parameter threshold: identity% −80, query coverage% −80, and e-value – 10^-5^.

Further to check if the horizontally acquired ARGs in phages were from the host chromosome or plasmid we used CARD prevalence data and further confirmed by mapping all ARGs sequences from CARD to the plasmid database (PLSDB: https://ccb-microbe.cs.uni-saarland.de/plsdb) [83] using BLASTN (identity% >= 95). The sources of ARGs carried by phages were found using the CARD prevalence data.

## Acknowledgments

Research in KK lab is supported by the Department of Science and Technology, DST [Indo-RSF, TPN-91844), Council of Scientific and Industrial Research (CSIR, 37/1752/23/EMR-II) Department of Biotechnology, DBT [BT/PR35928/BRB/10/1841/2019], and The Board of Research in Nuclear Sciences, BRNS [58/14/11/2020-BRNS/37188] grants. P. K. acknowledges financial support provided by the IIT Bombay Senior Research Fellowship.

## Author Contributions

**PK**: Conceptualization, Methodology, Formal analysis, Writing-Original draft preparation, Writing-Review & Editing; **BP:** Methodology, Investigation, Resources, Formal analysis, Writing-Editing, **TSP**: Methodology, Investigation, Resources, Formal Analysis, Writing-Editing, **KK:** Conceptualization, Methodology, Formal analysis, Writing-Original draft preparation; Writing-Review & Editing, Supervision, Funding acquisition.

