## Supplemental for "The low abundance of antimicrobial resistance genes (ARGs) in bacteriophages and their transfer bottlenecks limit the ability of phages to contribute to the spread of ARGs"

Supplementary Table S1. Estimation of ARG frequency in different databases

| Database | Total Phage/Prophage Sequences | No. of ARGs | Total no. of genes in the database | ARGs/Estimated all genes |
|---|---|---|---|---|
| NCBI | 38861 | 314 | 2466040 | $1.27 \times 10^{-4}$ |
| RefSeq | 5067 | 7 | 510990 | $1.37 \times 10^{-5}$ |
| IMGVR | 5576198 | 1360 | 111301861 | $1.22 \times 10^{-5}$ |
| GPD | 142809 | 169 | 7616074 | $2.22 \times 10^{-5}$ |
| CHVD | 44935 | 70 | 1940267 | $3.61 \times 10^{-5}$ |
| MGV | 189680 | 33 | 11887023 | $2.78 \times 10^{-6}$ |
| IGVD | 10021 | 4 | 342278 | $1.17 \times 10^{-5}$ |
| GOV2 | 195699 | 1 | 5017441 | $1.99 \times 10^{-7}$ |
| STV | 4065 | 0 | 149933 | 0 |
| Temperate Phage (TemPhD) | 66823 | 1815 | 7939937 | $2.29 \times 10^{-4}$ |
| Prophage (PhageClouds) | 49254 | 580 | 1298184 | $4.47 \times 10^{-4}$ |

Supplementary Table S2. ARG abundance in different bacterial genera.

| ARGs in Bacteria | | |
|---|---|---|
| Bacterial Genus | Number of ARG in bacterial genus | % of ARG in bacterial genus |
| Klebsiella | 2382 | 11.9 |
| Enterobacter | 1893 | 9.4 |
| Acinetobacter | 1850 | 9.2 |
| Pseudomonas | 1621 | 8.1 |
| Citrobacter | 1318 | 6.6 |
| Escherichia | 1189 | 5.9 |
| Shigella | 883 | 4.4 |
| Staphylococcus | 794 | 4.0 |
| Streptococcus | 599 | 3.0 |
| Vibrio | 587 | 2.9 |
| Aeromonas | 540 | 2.7 |
| Proteus | 477 | 2.4 |
| Providencia | 472 | 2.4 |
| Salmonella | 463 | 2.3 |
| Enterococcus | 391 | 2.0 |
| Serratia | 318 | 1.6 |
| Bacillus | 256 | 1.3 |
| Campylobacter | 241 | 1.2 |
| Morganella | 222 | 1.1 |
| Cronobacter | 205 | 1.0 |
| Burkholderia | 192 | 1.0 |
| Bacteroides | 165 | 0.8 |
| Raoultella | 128 | 0.6 |
| Elizabethkingia | 125 | 0.6 |
| Leclercia | 114 | 0.6 |
| Mycobacterium | 99 | 0.5 |
| Phocaeicola | 90 | 0.4 |
| Yersinia | 89 | 0.4 |
| Corynebacterium | 87 | 0.4 |
| Clostridium | 87 | 0.4 |
| Neisseria | 85 | 0.4 |
| Achromobacter | 82 | 0.4 |
| Stenotrophomonas | 71 | 0.4 |
| Listeria | 70 | 0.3 |
| Alcaligenes | 67 | 0.3 |
| Actinobacillus | 61 | 0.3 |
| Pasteurella | 59 | 0.3 |
| Laribacter | 59 | 0.3 |
| Bifidobacterium | 59 | 0.3 |
| Bordetella | 51 | 0.3 |
| Comamonas | 47 | 0.2 |
| Shewanella | 46 | 0.2 |
| Brevibacillus | 44 | 0.2 |
| Haemophilus | 43 | 0.2 |

| | | |
|---|---|---|
| Helicobacter | 37 | 0.2 |
| Enterocloster | 34 | 0.2 |
| Avibacterium | 34 | 0.2 |
| Moraxella | 32 | 0.2 |
| Parabacteroides | 32 | 0.2 |
| Brucella | 32 | 0.2 |
| Glaesserella | 31 | 0.2 |
| Ralstonia | 31 | 0.2 |
| Edwardsiella | 30 | 0.1 |
| Faecalibacterium | 30 | 0.1 |
| Myroides | 30 | 0.1 |
| Macrococcus | 29 | 0.1 |
| Lactobacillus | 28 | 0.1 |
| Lactococcus | 27 | 0.1 |
| Legionella | 27 | 0.1 |
| Eubacterium | 25 | 0.1 |
| Erysipelatoclostridium | 25 | 0.1 |
| Ruthenibacterium | 24 | 0.1 |
| Riemerella | 23 | 0.1 |
| Nocardia | 23 | 0.1 |
| Trueperella | 22 | 0.1 |
| Mycolicibacterium | 22 | 0.1 |
| Fusobacterium | 21 | 0.1 |
| Jeotgalibaca | 20 | 0.1 |
| Anaerostipes | 20 | 0.1 |
| Histophilus | 19 | 0.1 |
| Aliarcobacter | 18 | 0.1 |
| Prevotella | 18 | 0.1 |
| Kosakonia | 18 | 0.1 |
| Delftia | 16 | 0.1 |
| Sphingobium | 16 | 0.1 |
| Roseburia | 15 | 0.1 |
| Erysipelothrix | 14 | 0.1 |
| Xanthomonas | 14 | 0.1 |
| Plesiomonas | 13 | 0.1 |
| Capnocytophaga | 12 | 0.1 |
| Chryseobacterium | 12 | 0.1 |
| Leptospira | 12 | 0.1 |
| Blautia | 11 | 0.1 |
| Chlamydia | 11 | 0.1 |
| Clostridioides | 11 | 0.1 |
| Ligilactobacillus | 11 | 0.1 |
| Eggerthella | 11 | 0.1 |
| Rhodococcus | 11 | 0.1 |
| Rhizobium | 11 | 0.1 |
| Micrococcus | 10 | 0.0 |
| Parvimonas | 10 | 0.0 |

| | | |
|---|---|---|
| Francisella | 10 | 0.0 |
| Gemella | 9 | 0.0 |
| Paeniclostridium | 9 | 0.0 |
| Dysosmobacter | 9 | 0.0 |
| Treponema | 9 | 0.0 |
| Pectobacterium | 9 | 0.0 |
| Brachyspira | 9 | 0.0 |
| Kingella | 8 | 0.0 |
| Megasphaera | 8 | 0.0 |
| Mycoplasma | 8 | 0.0 |
| Collinsella | 8 | 0.0 |
| Christensenella | 8 | 0.0 |
| Lachnospiraceae | 8 | 0.0 |
| Herbinix | 8 | 0.0 |
| Sphingobacterium | 8 | 0.0 |
| Photorhabdus | 8 | 0.0 |
| Butyricimonas | 7 | 0.0 |
| Alistipes | 7 | 0.0 |
| Cutibacterium | 7 | 0.0 |
| Paenibacillus | 7 | 0.0 |
| Leminorella | 7 | 0.0 |
| Borrelia | 7 | 0.0 |
| Deinococcus | 6 | 0.0 |
| Gardnerella | 6 | 0.0 |
| Peptoniphilus | 6 | 0.0 |
| Gordonibacter | 6 | 0.0 |
| Mobiluncus | 6 | 0.0 |
| Propionibacterium | 6 | 0.0 |
| Borreliella | 6 | 0.0 |
| Massilistercora | 5 | 0.0 |
| Rothia | 5 | 0.0 |
| Peptacetobacter | 5 | 0.0 |
| Paracoccus | 5 | 0.0 |
| Rhodopseudomonas | 5 | 0.0 |
| Cupriavidus | 5 | 0.0 |
| Lysobacter | 5 | 0.0 |
| Glutamicibacter | 5 | 0.0 |
| Kocuria | 5 | 0.0 |
| Porphyromonas | 4 | 0.0 |
| Lachnoclostridium | 4 | 0.0 |
| Schaalia | 4 | 0.0 |
| Rhodanobacter | 4 | 0.0 |
| Aggregatibacter | 4 | 0.0 |
| Anaerococcus | 3 | 0.0 |
| Phoenicibacter | 3 | 0.0 |
| Inquilinus | 3 | 0.0 |
| Photobacterium | 3 | 0.0 |

| | | |
|---|---|---|
| Coxiella | 3 | 0.0 |
| Tannerella | 2 | 0.0 |
| Clostridiales | 2 | 0.0 |
| Luteibacter | 2 | 0.0 |
| Orrella | 2 | 0.0 |
| Pannonibacter | 2 | 0.0 |
| Nitrosomonas | 2 | 0.0 |
| Cytophaga | 2 | 0.0 |
| Alloiococcus | 2 | 0.0 |
| Eikenella | 2 | 0.0 |
| Parascardovia | 2 | 0.0 |
| Rhodobacteraceae | 1 | 0.0 |
| Muribaculaceae | 1 | 0.0 |
| Victivallales | 1 | 0.0 |
| Atopobiaceae | 1 | 0.0 |
| Methylorubrum | 1 | 0.0 |
| Thalassospira | 1 | 0.0 |
| Granulibacter | 1 | 0.0 |
| Dechloromonas | 1 | 0.0 |
| Anaplasma | 1 | 0.0 |
| Bartonella | 1 | 0.0 |
| Leifsonia | 1 | 0.0 |
| Cryptobacterium | 1 | 0.0 |
| Lancefieldella | 1 | 0.0 |
| Pseudoleptotrichia | 1 | 0.0 |
| Tropheryma | 1 | 0.0 |
| Actinomyces | 1 | 0.0 |
| Orientia | 1 | 0.0 |

Supplementary Table S3. Phage abundance in different bacterial genera.

| Phages in Bacteria | | |
|---|---|---|
| Bacterial Genus | Number of Phage in bacterial genus | % of Phage in bacterial genus |
| Mycolicibacterium | 675 | 13.0 |
| Escherichia | 517 | 9.9 |
| Pseudomonas | 331 | 6.4 |
| Salmonella | 295 | 5.7 |
| Gordonia | 273 | 5.3 |
| Staphylococcus | 184 | 3.5 |
| Lactococcus | 182 | 3.5 |
| Streptococcus | 180 | 3.5 |
| Klebsiella | 177 | 3.4 |
| Bacillus | 161 | 3.1 |
| Vibrio | 142 | 2.7 |
| Arthrobacter | 137 | 2.6 |
| Streptomyces | 121 | 2.3 |
| Microbacterium | 99 | 1.9 |
| Synechococcus | 85 | 1.6 |
| Mycobacterium | 75 | 1.4 |
| Cutibacterium | 75 | 1.4 |
| Erwinia | 65 | 1.3 |
| Acinetobacter | 65 | 1.3 |
| Enterococcus | 58 | 1.1 |
| Aeromonas | 55 | 1.1 |
| Shigella | 55 | 1.1 |
| Ralstonia | 54 | 1.0 |
| Yersinia | 45 | 0.9 |
| Burkholderia | 42 | 0.8 |
| Flavobacterium | 42 | 0.8 |
| Xanthomonas | 41 | 0.8 |
| Pectobacterium | 40 | 0.8 |
| Clostridium | 35 | 0.7 |
| Listeria | 34 | 0.7 |
| Paenibacillus | 31 | 0.6 |
| Stenotrophomonas | 29 | 0.6 |
| Serratia | 29 | 0.6 |
| Rhizobium | 28 | 0.5 |
| Cronobacter | 27 | 0.5 |
| Citrobacter | 26 | 0.5 |
| Rhodococcus | 24 | 0.5 |
| Cellulophaga | 23 | 0.4 |
| Lactobacillus | 23 | 0.4 |
| Proteus | 22 | 0.4 |
| Enterobacter | 20 | 0.4 |
| Pseudoalteromonas | 20 | 0.4 |
| Clostridioides | 19 | 0.4 |
| Prochlorococcus | 19 | 0.4 |

| | | |
|---|---|---|
| Enterobacteriaceae | 18 | 0.3 |
| Corynebacterium | 17 | 0.3 |
| Caulobacter | 17 | 0.3 |
| Leuconostoc | 17 | 0.3 |
| Lacticaseibacillus | 16 | 0.3 |
| Priestia | 16 | 0.3 |
| Achromobacter | 15 | 0.3 |
| Lactiplantibacillus | 15 | 0.3 |
| Dickeya | 12 | 0.2 |
| Propionibacterium | 11 | 0.2 |
| Agrobacterium | 10 | 0.2 |
| Shewanella | 9 | 0.2 |
| Pantoea | 9 | 0.2 |
| Providencia | 9 | 0.2 |
| Edwardsiella | 9 | 0.2 |
| Parasynechococcus | 8 | 0.2 |
| Sinorhizobium | 8 | 0.2 |
| Ruegeria | 8 | 0.2 |
| Faecalibacterium | 8 | 0.2 |
| Thermus | 8 | 0.2 |
| Campylobacter | 8 | 0.2 |
| Candidatus | 7 | 0.1 |
| Mannheimia | 7 | 0.1 |
| Rhodobacter | 7 | 0.1 |
| Brevibacillus | 7 | 0.1 |
| Alteromonas | 5 | 0.1 |
| Brucella | 5 | 0.1 |
| Polaribacter | 5 | 0.1 |
| Geobacillus | 5 | 0.1 |
| Bacteroides | 5 | 0.1 |
| Levilactobacillus | 5 | 0.1 |
| Rheinheimera | 5 | 0.1 |
| Xylella | 5 | 0.1 |
| Salinibacter | 5 | 0.1 |
| Tsukamurella | 5 | 0.1 |
| Chlamydia | 5 | 0.1 |
| Sulfitobacter | 5 | 0.1 |
| Spiroplasma | 5 | 0.1 |
| Dinoroseobacter | 4 | 0.1 |
| Roseobacter | 4 | 0.1 |
| Helicobacter | 4 | 0.1 |
| Bifidobacterium | 3 | 0.1 |
| Sphingomonas | 3 | 0.1 |
| Paeniglutamicibacter | 3 | 0.1 |
| Kosakonia | 3 | 0.1 |
| Brevibacterium | 3 | 0.1 |
| Tenacibaculum | 3 | 0.1 |

| | | |
|---|---|---|
| Leptospira | 3 | 0.1 |
| Nodularia | 3 | 0.1 |
| Marinomonas | 3 | 0.1 |
| Pasteurella | 3 | 0.1 |
| Phormidium | 3 | 0.1 |
| Delftia | 3 | 0.1 |
| Morganella | 3 | 0.1 |
| Limosilactobacillus | 3 | 0.1 |
| Mycoplasma | 3 | 0.1 |
| Oenococcus | 3 | 0.1 |
| Bdellovibrio | 3 | 0.1 |
| Brochothrix | 3 | 0.1 |
| Sutcliffiella | 2 | 0.0 |
| Curtobacterium | 2 | 0.0 |
| Carnobacterium | 2 | 0.0 |
| Maribacter | 2 | 0.0 |
| Mycobacteroides | 2 | 0.0 |
| Alkalihalobacillus | 2 | 0.0 |
| Raoultella | 2 | 0.0 |
| Lentibacter | 2 | 0.0 |
| Bordetella | 2 | 0.0 |
| Salinivibrio | 2 | 0.0 |
| Paracoccus | 2 | 0.0 |
| Idiomarinaceae | 2 | 0.0 |
| Weissella | 2 | 0.0 |
| Rhodovulum | 2 | 0.0 |
| Aggregatibacter | 2 | 0.0 |
| Parabacteroides | 2 | 0.0 |
| Psychrobacter | 2 | 0.0 |
| Croceibacter | 2 | 0.0 |
| Clavibacter | 2 | 0.0 |
| Leptolyngbya | 2 | 0.0 |
| Liberibacter | 2 | 0.0 |
| Nonlabens | 2 | 0.0 |
| Sodalis | 2 | 0.0 |
| Acholeplasma | 2 | 0.0 |
| Haemophilus | 2 | 0.0 |
| Glutamicibacter | 1 | 0.0 |
| Rothia | 1 | 0.0 |
| Ochrobactrum | 1 | 0.0 |
| Alcaligenes | 1 | 0.0 |
| bacterium | 1 | 0.0 |
| Winogradskyella | 1 | 0.0 |
| Olleya | 1 | 0.0 |
| Fusobacterium | 1 | 0.0 |
| Roseovarius | 1 | 0.0 |
| Butyrivibrio | 1 | 0.0 |

| | | |
|---|---|---|
| Photobacterium | 1 | 0.0 |
| Anoxybacillus | 1 | 0.0 |
| Aeribacillus | 1 | 0.0 |
| Sinomonas | 1 | 0.0 |
| Curvibacter | 1 | 0.0 |
| Eggerthella | 1 | 0.0 |
| Leclercia | 1 | 0.0 |
| Alphaproteobacteria | 1 | 0.0 |
| Gluconobacter | 1 | 0.0 |
| Acidovorax | 1 | 0.0 |
| Sphingobium | 1 | 0.0 |
| Aquamicrobium | 1 | 0.0 |
| Rhodoferax | 1 | 0.0 |
| Erysipelothrix | 1 | 0.0 |
| Puniceicoccales | 1 | 0.0 |
| Microcystis | 1 | 0.0 |
| Shimwellia | 1 | 0.0 |
| Hydrogenobaculum | 1 | 0.0 |
| [Brevibacterium] | 1 | 0.0 |
| Thermoanaerobacterium | 1 | 0.0 |
| Aurantimonas | 1 | 0.0 |
| Lelliottia | 1 | 0.0 |
| Mesorhizobium | 1 | 0.0 |
| Trichormus | 1 | 0.0 |
| Nitrincola | 1 | 0.0 |
| Aliivibrio | 1 | 0.0 |
| Loktanella | 1 | 0.0 |
| Salicola | 1 | 0.0 |
| Glaesserella | 1 | 0.0 |
| Tetrasphaera | 1 | 0.0 |
| Riemerella | 1 | 0.0 |
| Celeribacter | 1 | 0.0 |
| Colwellia | 1 | 0.0 |
| Salisaeta | 1 | 0.0 |
| Pediococcus | 1 | 0.0 |
| Nocardia | 1 | 0.0 |
| Planktothrix | 1 | 0.0 |
| Kluyvera | 1 | 0.0 |
| Halomonas | 1 | 0.0 |
| Iodobacter | 1 | 0.0 |
| Actinomyces | 1 | 0.0 |
| Thalassotalea | 1 | 0.0 |
| Azospirillum | 1 | 0.0 |
| Kitasatospora | 1 | 0.0 |
| alpha | 1 | 0.0 |
| Evansella | 1 | 0.0 |
| Myxococcus | 1 | 0.0 |

| | | |
|---|---|---|
| Rhodothermus | 1 | 0.0 |
| Actinoplanes | 1 | 0.0 |

Supplementary Table S4. Details of ARGs identified in different phage sequences.

| Phage | Phage Name | #ARGs | ARG Names | Phage Type | Completeness | #ARGs with MGE within 5k |
|---|---|---|---|---|---|---|
| MH445380.1 | Escherichia virus P1 | 11 | SHV-12,cmlA1,aadA2,mphA,dfrA14,ANT(3'')-Iia,qacL,TEM-34,sul3,Mrx,mef(B) | Temperate | complete | 11 |
| MN270279.1 | Streptococcus phage phi-SsuZKB4_rum | 7 | tet(40),mel,APH(3')-IIIa,ErmB,aad(6),SAT-4,tet(O) | Temperate | complete | 0 |
| BK046869.1 | Myoviridae sp. | 6 | vanG,vanT_in_vanG_cl,vanXY_in_vanG,vanW_in_vanG_cl,vanR_in_vanG_cl,vanU_in_vanG_cl | Temperate | partial | 0 |
| CAKLQF010000002.1 | Acinetobacter phage MD-2021a | 5 | adeG,adeF,adeL,adeH,abeS | Temperate | partial | 0 |
| CAKLQF020000002.1 | Acinetobacter phage MD-2021a | 5 | adeG,adeF,adeL,adeH,abeS | Temperate | partial | 0 |
| CAKLQH010000004.1 | Acinetobacter phage MD-2021a | 5 | adeG,adeF,adeL,adeH,abeS | Non-Temperate | partial | 0 |
| CAKLQH020000004.1 | Acinetobacter phage MD-2021a | 5 | adeG,adeF,adeL,adeH,abeS | Non-Temperate | partial | 0 |
| CP062450.1 | Staphylococcus phage ECel-2020c | 5 | msrA,APH(3')-IIIa,SAT-4,mphC,PC1_blaZ | Non-Temperate | complete | 0 |
| FN997652.1 | Streptococcus phage phi-SsUD.1 | 5 | APH(3')-IIIa,ErmB,aad(6),SAT-4,tet(W) | Non-Temperate | partial | 0 |
| KT336321.1 | Streptococcus phage phiSC070807 | 5 | mel,APH(3')-IIIa,ErmB,aad(6),SAT-4 | Temperate | complete | 0 |
| MN270273.1 | Streptococcus phage phi-SsuSC05017_rum | 5 | mel,APH(3')-IIIa,ErmB,aad(6),SAT-4 | Temperate | complete | 0 |
| CAKLQF010000015.1 | Acinetobacter phage MD-2021a | 4 | adeB,adeA,adeS,adeR | Temperate | partial | 0 |
| CAKLQF010000042.1 | Acinetobacter phage MD-2021a | 4 | sul1,arr-2,cmlA5,qacEdelta1 | Non-Temperate | partial | 4 |
| CAKLQF020000015.1 | Acinetobacter phage MD-2021a | 4 | adeB,adeA,adeS,adeR | Temperate | partial | 0 |
| CAKLQF020000042.1 | Acinetobacter phage MD-2021a | 4 | sul1,arr-2,cmlA5,qacEdelta1 | Non-Temperate | partial | 4 |
| CAKLQH010000024.1 | Acinetobacter phage MD-2021a | 4 | adeB,adeA,adeS,adeR | Temperate | partial | 0 |
| CAKLQH020000024.1 | Acinetobacter phage MD-2021a | 4 | adeB,adeA,adeS,adeR | Temperate | partial | 0 |

| | | | | | | |
|---|---|---|---|---|---|---|
| KX077896.1 | Streptococcus phage phiJH1301-2 | 4 | ANT(6)-Ia,mel,Lreu_cat-TC,AAC6_Ie_APH2_Ia | Temperate | complete | 1 |
| MK359990.1 | Streptococcus phage phi-SC181 | 4 | mel,optrA,Lreu_cat-TC,AAC6_Ie_APH2_Ia | Temperate | complete | 4 |
| MN270276.1 | Streptococcus phage phi-SsuSSJ28_rum | 4 | ANT(6)-Ia,ErmB,lnuD,AAC6_Ie_APH2_Ia | Temperate | complete | 3 |
| OQ572403.1 | Pseudomonas phage Y1 | 4 | MexJ,MexK,ArmR,MexL | Non-Temperate | complete | 0 |
| OQ572407.1 | Pseudomonas phage Y5 | 4 | arnA,mexP,mexQ,opmE | Non-Temperate | complete | 0 |
| CAKLQF010000003.1 | Acinetobacter phage MD-2021a | 3 | adeI,adeK,adeJ | Temperate | partial | 0 |
| CAKLQF010000043.1 | Acinetobacter phage MD-2021a | 3 | armA,mphE,msrE | Non-Temperate | partial | 3 |
| CAKLQF020000003.1 | Acinetobacter phage MD-2021a | 3 | adeI,adeK,adeJ | Temperate | partial | 0 |
| CAKLQF020000043.1 | Acinetobacter phage MD-2021a | 3 | armA,mphE,msrE | Non-Temperate | partial | 3 |
| CAKLQH010000002.1 | Acinetobacter phage MD-2021a | 3 | adeI,adeK,adeJ | Temperate | partial | 0 |
| CAKLQH010000068.1 | Acinetobacter phage MD-2021a | 3 | armA,mphE,msrE | Non-Temperate | partial | 3 |
| CAKLQH010000083.1 | Acinetobacter phage MD-2021a | 3 | arr-2,Sent_cmlA,cmlA5 | Non-Temperate | partial | 0 |
| CAKLQH020000002.1 | Acinetobacter phage MD-2021a | 3 | adeI,adeK,adeJ | Temperate | partial | 0 |
| CAKLQH020000068.1 | Acinetobacter phage MD-2021a | 3 | armA,mphE,msrE | Non-Temperate | partial | 3 |
| MN270277.1 | Streptococcus phage phi-SsuYTJ2_rum | 3 | tet(40),optrA,tet(O/W/32/O) | Temperate | partial | 1 |
| OP073502.1 | Bacteriophage sp. | 3 | dfrA15,qacEdelta1,sul1 | Non-Temperate | partial | 0 |
| BK020518.1 | Caudoviricetes sp | 2 | Eclo_acrA,acrB | Temperate | partial | 0 |
| CAKLQF010000005.1 | Acinetobacter phage MD-2021a | 2 | abeM,LpsB | Temperate | partial | 0 |
| CAKLQF010000037.1 | Acinetobacter phage MD-2021a | 2 | APH(6)-Id,APH(3)-Ib | Non-Temperate | partial | 0 |

| | | | | | | |
|---|---|---|---|---|---|---|
| CAKLQF020000005.1 | Acinetobacter phage MD-2021a | 2 | abeM,LpsB | Temperate | partial | 0 |
| CAKLQF020000037.1 | Acinetobacter phage MD-2021a | 2 | APH(6)-Id,APH(3)-Ib | Non-Temperate | partial | 0 |
| CAKLQH010000005.1 | Acinetobacter phage MD-2021a | 2 | abeM,LpsB | Temperate | partial | 0 |
| CAKLQH010000051.1 | Acinetobacter phage MD-2021a | 2 | APH(6)-Id,APH(3)-Ib | Non-Temperate | partial | 0 |
| CAKLQH010000071.1 | Acinetobacter phage MD-2021a | 2 | PER-7,sul1 | Non-Temperate | partial | 2 |
| CAKLQH020000005.1 | Acinetobacter phage MD-2021a | 2 | abeM,LpsB | Temperate | partial | 0 |
| CAKLQH020000071.1 | Acinetobacter phage MD-2021a | 2 | PER-7,sul1 | Non-Temperate | partial | 2 |
| CAKLQH020000083.1 | Acinetobacter phage MD-2021a | 2 | arr-2,cmlA5 | Non-Temperate | partial | 0 |
| FM864213.1 | Streptococcus phage phi-m46.1 | 2 | mel,tet(O) | Non-Temperate | partial | 0 |
| KF030445.1 | Escherichia phage 1720a-02 | 2 | APH(3')-IIIa,catA1 | Temperate | complete | 2 |
| KT336320.1 | Streptococcus phage phiNJ3 | 2 | mel,tet(O) | Temperate | complete | 1 |
| KT429160.1 | Staphylococcus phage SPbeta-like | 2 | dfrC,AAC6_Ie_APH2_Ia | Temperate | complete | 2 |
| MF172979.1 | Erysipelothrix phage phi1605 | 2 | mel,tet(M) | Temperate | complete | 0 |
| MN270258.1 | Streptococcus phage phi-SgaBSJ27_rum | 2 | tet(O),lnuC | Temperate | complete | 2 |
| MN270259.1 | Streptococcus phage phi-SgaBSJ31_rum | 2 | tet(O),lnuC | Temperate | complete | 2 |
| MN270261.1 | Streptococcus phage phi-SsuFJNP8_rum | 2 | tet(M),AAC6_Ie_APH2_Ia | Temperate | complete | 1 |
| MN270269.1 | Streptococcus phage phi-SsuHCJ3_rum | 2 | ANT(6)-Ia,AAC6_Ie_APH2_Ia | Temperate | complete | 2 |
| MT880872.1 | Staphylococcus phage PhiSepi-HH3 | 2 | dfrC,AAC6_Ie_APH2_Ia | Temperate | complete | 0 |
| NC_029119.1 | Staphylococcus phage SPbeta-like | 2 | dfrC,AAC6_Ie_APH2_Ia | Temperate | complete | 2 |
| MN270271.1 | Streptococcus phage phi-SsuNJ2_rum | 2 | mel,tet(O) | Temperate | complete | 0 |

| | | | | | | |
|---|---|---|---|---|---|---|
| MN270272.1 | Streptococcus phage phi-SsuNJ5_rum | 2 | mel,tet(O) | Temperate | complete | 0 |
| AF503408.1 | Enterobacteria phage P7 | 1 | TEM-34 | Temperate | complete | 1 |
| AY657002.1 | Streptococcus phage phi1207.3 | 1 | mel | Temperate | complete | 1 |
| BK019919.1 | Siphoviridae sp. | 1 | ErmG | Temperate | partial | 0 |
| BK026080.1 | Siphoviridae sp. | 1 | Lreu_cat-TC | Temperate | partial | 0 |
| BK026746.1 | Siphoviridae sp. | 1 | mel | Temperate | partial | 0 |
| BK027174.1 | Siphoviridae sp. | 1 | ErmG | Temperate | partial | 0 |
| BK027183.1 | Siphoviridae sp. | 1 | Ecol_emrE | Temperate | partial | 0 |
| BK027454.1 | Caudovirales sp. | 1 | *CfxA2* | Temperate | partial | 0 |
| BK029193.1 | Myoviridae sp. | 1 | *tetA(Q)* | Non-Temperate | partial | 0 |
| BK031860.1 | Caudoviricetes sp. | 1 | tet(O) | Temperate | partial | 1 |
| BK032296.1 | Caudoviricetes sp. | 1 | Ecol_emrE | Temperate | partial | 0 |
| BK041763.1 | Siphoviridae sp. | 1 | mel | Non-Temperate | partial | 0 |
| BK042198.1 | Siphoviridae sp. | 1 | lnuC | Temperate | partial | 1 |
| BK042658.1 | Myoviridae sp. | 1 | *tet(32)* | Non-Temperate | partial | 0 |
| BK044198.1 | Siphoviridae sp. | 1 | AAC6_Ie_APH2_Ia | Temperate | partial | 0 |
| BK044636.1 | Siphoviridae sp. | 1 | ACI-1 | Temperate | partial | 1 |
| BK044748.1 | Siphoviridae sp. | 1 | *mdtA* | Temperate | partial | 0 |
| BK047534.1 | Caudoviricetes sp | 1 | ACI-1 | Temperate | partial | 2 |
| BK047934.1 | Caudoviricetes sp. | 1 | catS | Non-Temperate | partial | 0 |
| BK048668.1 | Myoviridae sp. | 1 | *tet(W/N/W)* | Temperate | partial | 0 |
| BK052769.1 | Caudoviricetes sp. | 1 | ErmG | Temperate | partial | 1 |
| BK053916.1 | Myoviridae sp. | 1 | Ecol_emrE | Temperate | partial | 0 |
| BK055190.1 | Podoviridae sp. | 1 | Ecol_emrE | Temperate | partial | 0 |
| BK055476.1 | Siphoviridae sp. | 1 | lnuC | Temperate | partial | 0 |
| BK058939.1 | Bacteriophage sp. | 1 | Mef(En2) | Temperate | partial | 0 |
| CAKLQF010000001.1 | Acinetobacter phage MD-2021a | 1 | Abau_AbaF | Temperate | partial | 0 |
| CAKLQF010000006.1 | Acinetobacter phage MD-2021a | 1 | adeN | Temperate | partial | 0 |

| | | | | | | |
|---|---|---|---|---|---|---|
| CAKLQF010000008.1 | Acinetobacter phage MD-2021a | 1 | Abau_AbaQ | Non-Temperate | partial | 0 |
| CAKLQF010000009.1 | Acinetobacter phage MD-2021a | 1 | OXA-64 | Temperate | partial | 0 |
| CAKLQF010000018.1 | Acinetobacter phage MD-2021a | 1 | ADC-174 | Non-Temperate | partial | 0 |
| CAKLQF010000024.1 | Acinetobacter phage MD-2021a | 1 | Abau_AmvA | Temperate | partial | 0 |
| CAKLQF010000048.1 | Acinetobacter phage MD-2021a | 1 | OXA-23 | Non-Temperate | partial | 0 |
| CAKLQF010000052.1 | Acinetobacter phage MD-2021a | 1 | AAC(3)-IIe | Non-Temperate | partial | 1 |
| CAKLQF010000057.1 | Acinetobacter phage MD-2021a | 1 | AAC(6')-Ian | Non-Temperate | partial | 1 |
| CAKLQF010000063.1 | Acinetobacter phage MD-2021a | 1 | sul2 | Non-Temperate | partial | 0 |
| CAKLQF020000001.1 | Acinetobacter phage MD-2021a | 1 | Abau_AbaF | Temperate | partial | 0 |
| CAKLQF020000006.1 | Acinetobacter phage MD-2021a | 1 | adeN | Temperate | partial | 0 |
| CAKLQF020000008.1 | Acinetobacter phage MD-2021a | 1 | Abau_AbaQ | Non-Temperate | partial | 0 |
| CAKLQF020000009.1 | Acinetobacter phage MD-2021a | 1 | OXA-64 | Temperate | partial | 0 |
| CAKLQF020000018.1 | Acinetobacter phage MD-2021a | 1 | ADC-174 | Non-Temperate | partial | 0 |
| CAKLQF020000024.1 | Acinetobacter phage MD-2021a | 1 | Abau_AmvA | Temperate | partial | 0 |
| CAKLQF020000048.1 | Acinetobacter phage MD-2021a | 1 | OXA-23 | Non-Temperate | partial | 0 |
| CAKLQF020000052.1 | Acinetobacter phage MD-2021a | 1 | AAC(3)-IIe | Non-Temperate | partial | 1 |
| CAKLQF020000057.1 | Acinetobacter phage MD-2021a | 1 | AAC(6')-Ian | Non-Temperate | partial | 1 |
| CAKLQF020000063.1 | Acinetobacter phage MD-2021a | 1 | sul2 | Non-Temperate | partial | 0 |
| CAKLQH010000001.1 | Acinetobacter phage MD-2021a | 1 | Abau_AbaF | Temperate | partial | 0 |

| | | | | | | |
|---|---|---|---|---|---|---|
| CAKLQH010000012.1 | Acinetobacter phage MD-2021a | 1 | adeN | Temperate | partial | 0 |
| CAKLQH010000020.1 | Acinetobacter phage MD-2021a | 1 | Abau_AmvA | Temperate | partial | 0 |
| CAKLQH010000027.1 | Acinetobacter phage MD-2021a | 1 | Abau_AbaQ | Non-Temperate | partial | 0 |
| CAKLQH010000039.1 | Acinetobacter phage MD-2021a | 1 | OXA-64 | Temperate | partial | 0 |
| CAKLQH010000088.1 | Acinetobacter phage MD-2021a | 1 | OXA-23 | Non-Temperate | partial | 0 |
| CAKLQH010000093.1 | Acinetobacter phage MD-2021a | 1 | AAC(3)-IIe | Non-Temperate | partial | 1 |
| CAKLQH010000095.1 | Acinetobacter phage MD-2021a | 1 | AAC(6')-Ian | Non-Temperate | partial | 1 |
| CAKLQH010000100.1 | Acinetobacter phage MD-2021a | 1 | sul2 | Non-Temperate | partial | 0 |
| CAKLQH020000001.1 | Acinetobacter phage MD-2021a | 1 | Abau_AbaF | Temperate | partial | 0 |
| CAKLQH020000012.1 | Acinetobacter phage MD-2021a | 1 | adeN | Temperate | partial | 0 |
| CAKLQH020000020.1 | Acinetobacter phage MD-2021a | 1 | Abau_AmvA | Temperate | partial | 0 |
| CAKLQH020000027.1 | Acinetobacter phage MD-2021a | 1 | Abau_AbaQ | Non-Temperate | partial | 0 |
| CAKLQH020000039.1 | Acinetobacter phage MD-2021a | 1 | OXA-64 | Temperate | partial | 0 |
| CAKLQH020000051.1 | Acinetobacter phage MD-2021a | 1 | APH(6)-Id | Non-Temperate | partial | 0 |
| CAKLQH020000088.1 | Acinetobacter phage MD-2021a | 1 | OXA-23 | Non-Temperate | partial | 0 |
| CAKLQH020000093.1 | Acinetobacter phage MD-2021a | 1 | AAC(3)-IIe | Non-Temperate | partial | 1 |
| CAKLQH020000095.1 | Acinetobacter phage MD-2021a | 1 | AAC(6')-Ian | Non-Temperate | partial | 1 |
| CAKLQH020000100.1 | Acinetobacter phage MD-2021a | 1 | sul2 | Non-Temperate | partial | 0 |
| CP058330.1 | Klebsiella phage vB_Kpn_1825-KPC53 | 1 | KPC-25 | Non-Temperate | complete | 0 |

| | | | | | | |
|---|---|---|---|---|---|---|
| DQ222851.1 | Bacillus phage Cherry | 1 | FosB2 | Temperate | complete | 0 |
| DQ222853.1 | Bacillus phage Gamma | 1 | FosB2 | Temperate | complete | 0 |
| DQ289556.1 | Bacillus phage Gamma isolate d'Herelle | 1 | FosB2 | Temperate | complete | 0 |
| FO818745.1 | Escherichia phage RCS47 | 1 | SHV-2 | Temperate | complete | 1 |
| HM208303.1 | Escherichia phage vB_EcoP_24B | 1 | catA1 | Temperate | complete | 0 |
| KU238067.1 | Stx converting phage vB_EcoS_P27 | 1 | catA1 | Temperate | complete | 0 |
| KU238068.1 | Stx converting phage vB_EcoS_P32 | 1 | catA1 | Temperate | complete | 0 |
| KU238069.1 | Stx converting phage vB_EcoS_P22 | 1 | catA1 | Temperate | complete | 0 |
| KU238070.1 | Stx converting phage vB_EcoS_ST2-8624 | 1 | catA1 | Temperate | complete | 0 |
| KU760857.1 | Salmonella phage SJ46 | 1 | CTX-M-27 | Temperate | complete | 1 |
| KY065497.1 | Streptococcus phage IPP61 | 1 | tet(M) | Temperate | complete | 1 |
| KY304108.1 | Escherichia virus T4 | 1 | *CTX-M-108* | Non-Temperate | partial | 0 |
| KY304109.1 | Escherichia virus T4 | 1 | *CTX-M-108* | Non-Temperate | partial | 0 |
| KY785037.1 | Escherichia phage Jp6 | 1 | *CTX-M-108* | Non-Temperate | partial | 0 |
| LR595861.1 | Escherichia virus Lambda_4C10 | 1 | Ecol_emrE | Temperate | partial | 0 |
| LR595864.1 | Escherichia virus Lambda_4A7 | 1 | Ecol_emrE | Temperate | partial | 0 |
| M14017.1 | Enterobacteria phage f1 | 1 | TEM-68 | Non-Temperate | partial | 0 |
| MK448400.1 | Streptococcus satellite phage Javan285 | 1 | lnuC | Temperate | complete | 1 |
| MK448483.1 | Streptococcus satellite phage Javan415 | 1 | lnuC | Temperate | complete | 0 |
| MK448997.1 | Streptococcus phage Javan630 | 1 | ANT(6)-Ia | Temperate | complete | 0 |
| MN270263.1 | Streptococcus phage phi-SsuFJSM5_rum | 1 | ANT(6)-Ia | Temperate | complete | 0 |
| MN270264.1 | Streptococcus phage phi-SsuFJSM7_rum | 1 | ANT(6)-Ia | Temperate | complete | 0 |

| | | | | | | |
|---|---|---|---|---|---|---|
| MN270274.1 | Streptococcus phage phi-SsuSH0918_comEC | 1 | dfrG | Temperate | partial | 0 |
| MN399839.1 | uncultured phage | 1 | TEM-116 | Non-Temperate | partial | 0 |
| MN399840.1 | uncultured phage | 1 | TEM-116 | Non-Temperate | partial | 0 |
| MN399841.1 | uncultured phage | 1 | TEM-116 | Non-Temperate | partial | 0 |
| MN399842.1 | uncultured phage | 1 | TEM-116 | Non-Temperate | partial | 0 |
| MN399843.1 | uncultured phage | 1 | TEM-116 | Non-Temperate | partial | 0 |
| MN399844.1 | uncultured phage | 1 | TEM-116 | Non-Temperate | partial | 0 |
| MN399845.1 | uncultured phage | 1 | TEM-116 | Non-Temperate | partial | 0 |
| MN399846.1 | uncultured phage | 1 | TEM-116 | Non-Temperate | partial | 0 |
| MN399847.1 | uncultured phage | 1 | TEM-116 | Non-Temperate | partial | 0 |
| MN399848.1 | uncultured phage | 1 | TEM-116 | Non-Temperate | partial | 0 |
| MN399849.1 | uncultured phage | 1 | TEM-116 | Non-Temperate | partial | 0 |
| MN399850.1 | uncultured phage | 1 | TEM-116 | Non-Temperate | partial | 0 |
| MN399851.1 | uncultured phage | 1 | TEM-229 | Non-Temperate | partial | 0 |
| MN688132.1 | Enterobacter phage LAU1 | 1 | CTX-M-15 | Non-Temperate | complete | 1 |
| MT303952.1 | Streptococcus phage phi29862 | 1 | mel | Temperate | complete | 1 |
| MT311967.1 | Streptococcus phage phi29854 | 1 | mel | Temperate | complete | 1 |
| MT311968.1 | Streptococcus phage phi29961 | 1 | mel | Temperate | complete | 1 |
| MT835479.1 | Bacteriophage sp. | 1 | mphO | Non-Temperate | partial | 0 |
| MZ375336.1 | Salmonella phage seszw | 1 | catA1 | Non-Temperate | complete | 0 |

| | | | | | | |
|---|---|---|---|---|---|---|
| NC_007458.1 | Bacillus phage Gamma | 1 | FosB2 | Temperate | complete | 0 |
| NC_027984.1 | Escherichia phage vB_EcoP_24B | 1 | catA1 | Temperate | complete | 0 |
| NC_031129.1 | Salmonella phage SJ46 | 1 | CTX-M-27 | Temperate | complete | 1 |
| NC_042128.1 | Escherichia phage RCS47 | 1 | SHV-2 | Temperate | complete | 1 |
| NC_049922.1 | Stx converting phage vB_EcoS_ST2-8624 | 1 | catA1 | Temperate | complete | 0 |
| NC_049925.1 | Stx converting phage vB_EcoS_P27 | 1 | catA1 | Temperate | complete | 0 |
| NC_049946.1 | Escherichia virus Lambda_4A7 | 1 | Ecol_emrE | Temperate | complete | 0 |
| NC_050152.1 | Enterobacteria phage P7 | 1 | TEM-34 | Temperate | complete | 1 |
| OM475428.1 | Escherichia phage 1W | 1 | Ecol_emrE | Temperate | partial | 0 |
| OM475430.1 | Escherichia phage 3W | 1 | Ecol_emrE | Temperate | partial | 0 |
| ON018986.2 | Escherichia phage JL22 | 1 | CTX-M-55 | Temperate | complete | 1 |
| ON470620.1 | Escherichia phage vB_EcoM-813R9 | 1 | Ecol_mdfA | Temperate | partial | 0 |
| OP076392.1 | Bacteriophage sp. | 1 | ErmB | Temperate | partial | 0 |
| OP328421.1 | Staphylococcus phage phiSa2wa-st1420 | 1 | dfrG | Temperate | partial | 1 |
| OP328422.1 | Staphylococcus phage phiSa2wa-st1633 | 1 | dfrG | Temperate | complete | 1 |
| OR487170.1 | Bacillus phage FI_KG-Lek | 1 | FosB2 | Non-Temperate | complete | 0 |
| OV696612.1 | Escherichia phage Perceval | 1 | Ecol_emrE | Temperate | complete | 0 |
| Z35638.1 | Escherichia virus phiX174 | 1 | TEM-116 | Non-Temperate | partial | 0 |

Syupplementary Table S5. Host of ARGs carrying phages are also among the fast growing bacteria

| **Fast Growing Species as per Touchon et al. 2016** |
|---|
| Escherichia coli |
| Staphylococcus aureus |
| Salmonella enterica |
| Listeria monocytogenes |
| Streptococcus pneumoniae |
| Streptococcus pyogenes |
| Streptococcus suis |
| Bacillus cereus |
| Clostridium botulinum |
| Corynebacterium diphtheriae |
| Yersinia pestis |
| Acinetobacter baumannii |
| Bacillus amyloliquefaciens |
| Bifidobacterium longum |
| Haemophilus influenzae |
| Pseudomonas putida KT2440 |
| Acetobacter pasteurianus |
| Bacillus thuringiensis |
| Lactococcus lactis |
| Vibrio cholerae |
| Bacillus anthracis |
| Bacillus subtilis |
| Burkholderia pseudomallei |
| Pseudomonas aeruginosa |
| Shewanella baltica |
| Alteromonas macleodii |
| Klebsiella pneumoniae |
| Pseudomonas stutzeri |
| Streptococcus thermophilus |
| Enterobacter cloacae |
| Pseudomonas fluorescens |
| Ralstonia solanacearum |
| Riemerella anatipestifer |
| Streptococcus agalactiae |
| Brachyspira pilosicoli |
| Burkholderia cenocepacia AU 1054 |
| Clostridium difficile |
| Enterococcus faecalis |
| Pantoea ananatis |
| Pasteurella multocida |
| Rhodobacter sphaeroides |
| Shigella flexneri |
| Streptococcus dysgalactiae |
| Streptococcus equi |
| Stenotrophomonas maltophilia |

Comparison

| **Host of ARG carrying phage** |
|---|
| Acinetobacter |
| Streptococcus |
| Unknown |
| Escherichia |
| Staphylococcus |
| Pseudomonas |
| Bacillus |
| Enterobacteria |
| Salmonella |
| Erysipelothrix |
| Klebsiella |

**Host of all the ARG carrying phages are fast growing bacteria**

| |
|---|
| Yersinia pseudotuberculosis |
| Actinobacillus pleuropneumoniae |
| Aggregatibacter actinomycetemcomitans |
| Bacteroides fragilis |
| Bordetella bronchiseptica |
| Bordetella pertussis |
| Clostridium perfringens |
| Corynebacterium glutamicum |
| Corynebacterium ulcerans |
| Dickeya dadantii |
| Gardnerella vaginalis |
| Lactobacillus johnsonii |
| Lactobacillus plantarum |
| Lactobacillus reuteri |
| Neisseria gonorrhoeae |
| Paenibacillus mucilaginosus |
| Paenibacillus polymyxa |
| Pseudomonas syringae |
| Streptococcus gallolyticus |
| Streptococcus salivarius |
| Vibrio vulnificus |
| Yersinia enterocolitica |
| Arcobacter butzleri |
| Bacillus licheniformis |
| Bdellovibrio bacteriovorus |
| Bordetella parapertussis |
| Burkholderia ambifaria |
| Burkholderia multivorans |
| Edwardsiella tarda |
| Enterococcus faecium |
| Erwinia pyrifoliae |
| Gluconobacter oxydans |
| Haemophilus somnus |
| Hydrogenobacter thermophilus |
| Ketogulonicigenium vulgare |
| Klebsiella oxytoca |
| Lactobacillus acidophilus |
| Lactobacillus amylovorus |
| Lactobacillus fermentum |
| Lactococcus garvieae |
| Lactobacillus helveticus H10 |
| Lactobacillus salivarius |
| Leuconostoc mesenteroides |
| Pectobacterium carotovorum |
| Phaeobacter gallaeciensis |
| Rahnella aquatilis |
| Ralstonia eutropha |

| |
|---|
| Ralstonia pickettii |
| Shigella boydii |
| Shewanella oneidensis |
| Shewanella putrefaciens |
| Shigella sonnei |
| Shewanella sp. MR-7 |
| Sinorhizobium fredii HH103 |
| Streptomyces cattleya |
| Staphylococcus epidermidis |
| Staphylococcus lugdunensis |
| Staphylococcus pseudintermedius |
| Synechococcus elongatus |
| Variovorax paradoxus |
| Vibrio fischeri |
| Vibrio parahaemolyticus |
| Acinetobacter oleivorans |
| Acidaminococcus fermentans |
| Acidaminococcus intestini |
| Acidovorax sp. KKS102 |
| Acholeplasma laidlawii |
| Actinoplanes missouriensis |
| Actinoplanes sp. |
| Actinobacillus suis |
| Acinetobacter sp. ADP1 |
| Actinobacillus succinogenes |
| Achromobacter xylosoxidans |
| Advenella kashmirensis |
| Aeromonas salmonicida |
| Aeromonas veronii |
| Aggregatibacter aphrophilus |
| Agrobacterium sp. H13-3 |
| Agrobacterium radiobacter |
| Agrobacterium tumefaciens |
| Alcanivorax dieselolei |
| Alkaliphilus metalliredigens |
| Aliivibrio salmonicida |
| Alteromonas sp. SN2 |
| Amphibacillus xylanus |
| Anoxybacillus flavithermus |
| Anaerococcus prevotii |
| Caldicellulosiruptor bescii |
| Arcanobacterium haemolyticum |
| Arcobacter sp. 90 |
| Asticcacaulis excentricus |
| Atopobium parvulum |
| Azotobacter vinelandii |
| Bacillus atrophaeus |

| |
|---|
| Bacillus clausii |
| Bacillus cellulosilyticus |
| Bacillus halodurans |
| Bacillus sp. JS |
| Bacteriovorax marinus |
| Bacillus pumilus |
| Bacillus selenitireducens |
| Bacillus weihenstephanensis |
| Bordetella avium |
| Bordetella petrii |
| Brevibacillus brevis |
| Brachyspira intermedia |
| Brachyspira murdochii |
| Burkholderia sp. 383 |
| Burkholderia cepacia |
| Burkholderia sp. CCGE1002 |
| Burkholderia gladioli |
| Burkholderia sp. CCGE1001 |
| Burkholderia glumae |
| Burkholderia sp. KJ006 |
| Burkholderia phymatum |
| Burkholderia phytofirmans |
| Burkholderia thailandensis |
| Burkholderia vietnamiensis |
| Burkholderia xenovorans |
| Burkholderia sp. YI23 |
| Carnobacterium sp. |
| Catenulispora acidiphila |
| Caldicellulosiruptor hydrothermalis |
| Carnobacterium maltaromaticum |
| Caldicellulosiruptor owensensis |
| Cellulomonas fimi |
| Cellvibrio gilvus |
| Cellvibrio japonicus |
| Chitinophaga pinensis |
| Chromohalobacter salexigens |
| Chromobacterium violaceum |
| Citrobacter koseri |
| Citrobacter rodentium |
| Clostridium acidurici |
| Clostridium beijerinckii |
| Clostridium cellulovorans |
| Clostridium lentocellum |
| Clostridium novyi |
| Clostridium phytofermentans |
| Clostridium saccharolyticum |
| Clostridium sticklandii |

| |
|---|
| Clostridium sp. SY8519 |
| Clostridium tetani |
| Corynebacterium aurimucosum |
| Corynebacterium efficiens |
| Collimonas fungivorans |
| Corynebacterium kroppenstedtii |
| Corynebacterium resistens |
| Comamonas testosteroni |
| Corynebacterium urealyticum |
| Corynebacterium variabile |
| Cronobacter sakazakii ES15 |
| Cronobacter turicensis |
| Cupriavidus necator |
| Cupriavidus taiwanensis |
| Delftia acidovorans |
| Delftia sp. Cs1-4 |
| Deinococcus gobiensis |
| Deinococcus maricopensis |
| Deinococcus peraridilitoris |
| Desulfobulbus propionicus |
| Deinococcus radiodurans |
| Desulfotomaculum reducens |
| Desulfovibrio salexigens |
| Deinococcus deserti |
| Dechlorosoma suillum |
| Dickeya zeae |
| Dyadobacter fermentans |
| Echinicola vietnamensis |
| Edwardsiella ictaluri |
| Enterobacter aerogenes |
| Enterobacter asburiae |
| Enterobacteriaceae bacterium |
| Enterococcus hirae |
| Cronobacter sakazakii ATCC BAA-894 |
| Enterobacter sp. |
| Erwinia billingiae |
| Pectobacterium atrosepticum |
| Erwinia sp. Ejp617 |
| Erysipelothrix rhusiopathiae |
| Erwinia tasmaniensis |
| Escherichia blattae |
| Escherichia fergusonii |
| Eubacterium limosum |
| Eubacterium rectale |
| Exiguobacterium sp. |
| Finegoldia magna |
| Frateuria aurantia |

| |
|---|
| Gallibacterium anatis |
| Geobacillus kaustophilus |
| Geobacillus sp. C56-T3 |
| Paenibacillus sp. Y412MC10 |
| Geobacillus sp. Y412MC61 |
| Geobacillus thermoglucosidasius |
| Geobacillus thermodenitrificans |
| Geobacillus thermoleovorans |
| Geobacillus sp. Y412MC52 |
| Geobacillus sp. Y4.1MC1 |
| Gordonia sp. KTR9 |
| Gramella forsetii |
| Hahella chejuensis |
| Haemophilus ducreyi |
| Halomonas elongata |
| Haemophilus parasuis |
| Haemophilus parainfluenzae |
| Herpetosiphon aurantiacus |
| Herbaspirillum seropedicae |
| Idiomarina loihiensis |
| Ilyobacter polytropus |
| Janthinobacterium sp. |
| Jonesia denitrificans |
| Kineococcus radiotolerans |
| Klebsiella variicola |
| Lactobacillus brevis |
| Lactobacillus crispatus |
| Lactobacillus gasseri |
| Laribacter hongkongensis |
| Lactobacillus ruminis |
| Leuconostoc sp. C2 |
| Leuconostoc carnosum |
| Leuconostoc citreum |
| Leuconostoc gasicomitatum |
| Leuconostoc kimchii |
| Listeria innocua |
| Listeria seeligeri |
| Listeria welshimeri |
| Lysinibacillus sphaericus |
| Marinobacter adhaerens |
| Macrococcus caseolyticus |
| Marinomonas mediterranea |
| Marinomonas sp. MWYL1 |
| Marinitoga piezophila |
| Mannheimia succiniciproducens |
| Megasphaera elsdenii |
| Melissococcus plutonius ATCC 35311 |

| |
|---|
| Melissococcus plutonius DAT561 |
| Methylovorus glucosetrophus |
| Meiothermus silvanus |
| Micavibrio aeruginosavorus |
| Mycobacterium abscessus |
| Neisseria lactamica |
| Nitratifractor salsuginis |
| Novosphingobium sp. |
| Novosphingobium aromaticivorans |
| Nocardioides sp. |
| Nocardia farcinica |
| Ochrobactrum anthropi |
| Oceanimonas sp. GK1 |
| Pantoea sp. At-9b |
| Parabacteroides distasonis |
| Paenibacillus sp. JDR-2 |
| Paenibacillus terrae |
| Pantoea vagans |
| Pediococcus claussenii |
| Persephonella marina |
| Pediococcus pentosaceus |
| Pedobacter saltans |
| Pectobacterium wasabiae |
| Photorhabdus asymbiotica |
| Photorhabdus luminescens |
| Photobacterium profundum |
| Prevotella intermedia |
| Proteus mirabilis |
| Providencia stuartii |
| Pseudoalteromonas atlantica |
| Pseudomonas brassicacearum |
| Pseudonocardia dioxanivorans |
| Pseudomonas entomophila |
| Pseudovibrio sp. FO-BEG1 |
| Pseudomonas fulva |
| Pseudoalteromonas haloplanktis |
| Pseudomonas mendocina NK-01 |
| Pseudomonas mendocina ymp |
| Pseudogulbenkiania sp. |
| Rahnella sp. Y9602 |
| Rhodobacter capsulatus |
| Rhodococcus erythropolis |
| Rhodococcus opacus |
| Rhodococcus jostii |
| Rhizobium tropici |
| Salinispora arenicola |
| Salmonella bongori |

| |
|---|
| Saccharothrix espanaensis |
| Salinispora tropica |
| Serratia sp. AS9 |
| Serratia sp. AS12 |
| Serratia sp. AS13 |
| Serratia marcescens |
| Serratia proteamaculans |
| Selenomonas ruminantium |
| Shewanella denitrificans |
| Shigella dysenteriae |
| Shewanella frigidimarina |
| Shewanella halifaxensis |
| Sinorhizobium medicae |
| Ruegeria sp. TM1040 |
| Solitalea canadensis |
| Sodalis glossinidius |
| Solibacillus silvestris |
| Streptomyces avermitilis |
| Streptomyces bingchenggensis |
| Staphylococcus carnosus |
| Staphylococcus haemolyticus |
| Streptococcus infantarius |
| Streptococcus mitis |
| Streptococcus parauberis |
| Streptococcus pseudopneumoniae |
| Staphylococcus saprophyticus |
| Streptomyces scabiei |
| Streptomyces venezuelae |
| Taylorella asinigenitalis |
| Tetragenococcus halophilus |
| Teredinibacter turnerae |
| Thermovibrio ammonificans |
| Thermosipho melanesiensis |
| Thermaerobacter marianensis |
| Thermosediminibacter oceani |
| Thioalkalivibrio sulfidophilus |
| Tolumonas auensis |
| Tsukamurella paurometabola |
| Vibrio sp. EJY3 |
| Vibrio furnissii |
| Vibrio harveyi |
| Weissella koreensis |
| Xenorhabdus bovienii |
| Xenorhabdus nematophila |

Supplementary Table S6. Using online BLASTN for identifying Streptococcus phage ARG cluster by using it as query in host bacteria

| Description | Max Score | Total Score | Query Coverage | E value | Identity% | Accession |
|---|---|---|---|---|---|---|
| *Streptococcus suis strain 1135-10* | 32297 | 37180 | 98% | 0 | 94.44 | KY400496.1 |
| *Streptococcus suis isolate 9401240 genome assembly, chromosome: 1* | 29933 | 37017 | 95% | 0 | 95.64 | LR738724.1 |
| *Streptococcus suis strain M106471_S40 chromosome, complete genome* | 29529 | 73538 | 99% | 0 | 95.25 | CP102135.1 |
| *Streptococcus suis strain AKJ18 chromosome, complete genome* | 29403 | 36933 | 97% | 0 | 95.13 | CP082205.1 |
| *Streptococcus suis strain Transconjugant cAKJ18 chromosome, complete genome* | 29403 | 36933 | 97% | 0 | 95.13 | CP082201.1 |
| *Streptococcus suis strain 12RC1 chromosome, complete genome* | 28995 | 36566 | 95% | 0 | 95.17 | CP102094.1 |
| *Streptococcus suis strain Transconjugant cSS389 chromosome, complete genome* | 27409 | 36601 | 94% | 0 | 95.14 | CP082197.1 |
| *Streptococcus suis strain SS389 chromosome, complete genome* | 27409 | 36601 | 94% | 0 | 95.14 | CP082202.1 |
| *Streptococcus suis strain FJNP8 clone phi_SsuFJNP8_rum genomic sequence* | 26576 | 36714 | 92% | 0 | 95.85 | MZ960466.1 |
| *Streptococcus suis strain JSXZP1 clone phi_SsuJSXZP1_comEC genomic sequence* | 26441 | 36074 | 92% | 0 | 95.79 | MZ960473.1 |
| *Streptococcus suis strain XMSJ29 clone phi_SsuXMSJ29_comEC genomic sequence* | 26378 | 36011 | 92% | 0 | 95.71 | MZ960488.1 |
| *Streptococcus suis strain JSXZP2 clone phi_SsuJSXZP2_comEC genomic sequence* | 26374 | 36008 | 92% | 0 | 95.71 | MZ960475.1 |
| *Streptococcus suis strain XMJ29 clone phi_SsuXMJ29_comEC genomic sequence* | 26210 | 36679 | 95% | 0 | 95.53 | MZ960487.1 |
| *Streptococcus suis strain SHJ7 clone phi_SsuSHJ7_comEC genomic sequence* | 26210 | 36679 | 95% | 0 | 95.53 | MZ960485.1 |
| *Streptococcus dysgalactiae strain DY107 chromosome, complete genome* | 26083 | 35610 | 92% | 0 | 95.39 | CP082206.1 |
| *Streptococcus suis strain Transconjugant cDY107 chromosome, complete genome* | 26083 | 35610 | 92% | 0 | 95.39 | CP082200.1 |
| *Streptococcus suis strain CZ130302 chromosome, complete genome* | 25736 | 35579 | 92% | 0 | 95 | CP024974.1 |
| *Streptococcus suis strain GMJ14 clone phi_SsuGMJ14_rum genomic sequence* | 25736 | 35386 | 92% | 0 | 95.01 | MZ960470.1 |
| *Streptococcus suis strain GMJ13 clone phi_SsuGMJ13_rum genomic sequence* | 25736 | 35386 | 92% | 0 | 95.01 | MZ960469.1 |
| *Streptococcus suis strain GMB2P clone phi_SsuGMB2P_rum genomic sequence* | 25736 | 35386 | 92% | 0 | 95.01 | MZ960468.1 |
| *Streptococcus suis strain JS7 clone phi_SsuJS7_rum genomic sequence* | 25671 | 35437 | 92% | 0 | 94.94 | MZ960471.1 |
| *Streptococcus suis strain SC183 chromosome, complete genome* | 24068 | 34367 | 94% | 0 | 93.77 | CP071305.1 |
| *Streptococcus suis strain LS9N genome assembly, chromosome: I* | 22940 | 33025 | 87% | 0 | 94.73 | LT671674.1 |

| | | | | | | |
|---|---|---|---|---|---|---|
| *Streptococcus suis strain 110 clone phi_Ssu110_rum genomic sequence* | 22242 | 34103 | 91% | 0 | 93.88 | MZ960465.1 |
| *Streptococcus suis DAT300 DNA, complete genome* | 21664 | 35867 | 92% | 0 | 95.27 | AP023392.1 |
| *Streptococcus suis strain DNS20 chromosome, complete genome* | 21431 | 35107 | 92% | 0 | 94.96 | CP102145.1 |
| *Streptococcus acidominimus strain NCTC11291 genome assembly, chromosome: 1* | 21241 | 35933 | 95% | 0 | 94.68 | LT906454.1 |
| *Streptococcus suis strain XMSJ29 clone phi_SsuXMSJ29_rum genomic sequence* | 21211 | 35274 | 91% | 0 | 94.68 | MZ960489.1 |
| *Streptococcus oriscaviae strain HKU75 chromosome, complete genome* | 21178 | 32900 | 87% | 0 | 95.33 | CP073084.1 |
| *Streptococcus suis strain JSXZP2 clone phi_SsuJSXZP2_rum genomic sequence* | 21167 | 35229 | 91% | 0 | 94.62 | MZ960476.1 |
| *Streptococcus suis strain JSXZP1 clone phi_SsuJSXZP1_rum genomic sequence* | 21156 | 35218 | 91% | 0 | 94.6 | MZ960474.1 |
| *Streptococcus suis strain 1081 chromosome, complete genome* | 20620 | 34545 | 93% | 0 | 93.73 | CP017667.1 |
| *Streptococcus suis strain JSB1 clone phi_SsuJSB1_rum genomic sequence* | 20567 | 32693 | 87% | 0 | 94.89 | MZ960472.1 |
| *Streptococcus suis strain SHSJ7 clone phi_SsuSHSJ7_comEC genomic sequence* | 17335 | 35022 | 92% | 0 | 94.77 | MZ960486.1 |
| *Streptococcus suis strain CS100322 chromosome, complete genome* | 16899 | 33439 | 92% | 0 | 94.06 | CP024050.1 |
| *Streptococcus suis strain YY060816 composite mobile genetic element CMGEYY060816 mobile element, complete sequence* | 16899 | 33439 | 92% | 0 | 94.06 | KX077898.1 |
| *Streptococcus suis strain TZ080501 composite mobile genetic element CMGETZ080501 mobile element, complete sequence* | 16899 | 33439 | 92% | 0 | 94.06 | KX077897.1 |
| *Streptococcus suis strain SZ1908 chromosome, complete genome* | 16899 | 33433 | 92% | 0 | 94.06 | CP082948.1 |
| *Streptococcus suis SC070731, complete genome* | 16899 | 33439 | 92% | 0 | 94.06 | CP003922.1 |
| *Streptococcus suis JS14, complete genome* | 16862 | 33396 | 92% | 0 | 94 | CP002465.1 |
| *Streptococcus suis strain HA1003 chromosome, complete genome* | 16783 | 67857 | 92% | 0 | 93.87 | CP030125.1 |
| *Streptococcus suis strain YSB3 clone phi_SsuYSB3_rum genomic sequence* | 16768 | 34188 | 92% | 0 | 93.86 | MZ960490.1 |
| *Streptococcus suis strain YSJ17 chromosome, complete genome* | 16768 | 34193 | 92% | 0 | 93.86 | CP032064.1 |
| *Streptococcus suis strain MLYF4-9 clone phi_SsuMLYF4-9_rum genomic sequence* | 16652 | 33651 | 89% | 0 | 93.87 | MZ960484.1 |
| *Streptococcus suis strain MLBY4-9 clone phi_SsuMLBY4-9_rum genomic sequence* | 16652 | 33651 | 89% | 0 | 93.87 | MZ960483.1 |
| *Streptococcus suis strain MLBY4-8 clone phi_SsuMLBY4-8_rum genomic sequence* | 16652 | 33651 | 89% | 0 | 93.87 | MZ960482.1 |

| | | | | | | |
|---|---|---|---|---|---|---|
| *Streptococcus suis strain MLB3-3B clone phi_SsuMLB3-3B_rum genomic sequence* | 16652 | 33651 | 89% | 0 | 93.87 | MZ960481.1 |
| *Streptococcus suis strain ML3-36 clone phi_SsuML3-36_rum genomic sequence* | 16652 | 33651 | 89% | 0 | 93.87 | MZ960480.1 |
| *Streptococcus suis strain ML18-37d-1 clone phi_SsuML18-37d-1_rum genomic sequence* | 16652 | 33651 | 89% | 0 | 93.87 | MZ960479.1 |
| *Streptococcus suis strain ML18-100d-21 clone phi_SsuML18-100d-21_rum genomic sequence* | 16652 | 33651 | 89% | 0 | 93.87 | MZ960478.1 |
| *Streptococcus suis strain ML18-100d-20 clone phi_SsuML18-100d-20_rum genomic sequence* | 16652 | 33651 | 89% | 0 | 93.87 | MZ960477.1 |
| *Streptococcus dysgalactiae subsp. equisimilis AC-2713, complete genome* | 16517 | 54313 | 92% | 0 | 93.44 | HE858529.1 |
| *Streptococcus suis D12, complete genome* | 16295 | 33033 | 89% | 0 | 93.08 | CP002644.1 |
| *Streptococcus suis strain YTJ2 clone phi_SsuYTJ2_comEC genomic sequence* | 16246 | 32724 | 92% | 0 | 93.01 | MZ960491.1 |
| *Streptococcus suis strain SC215 lsa(E) multiresistance gene cluster, complete sequence* | 16222 | 32560 | 89% | 0 | 92.97 | MN437485.1 |
| *Streptococcus suis strain FJSM7 clone phi_SsuFJSM7_rum genomic sequence* | 15187 | 36308 | 95% | 0 | 94.46 | MZ960467.1 |
| *Streptococcus suis strain FJSM5 chromosome, complete genome* | 15187 | 36308 | 95% | 0 | 94.46 | CP082204.1 |
| *Streptococcus suis strain Transconjugant cFJSM5 chromosome, complete genome* | 15187 | 36308 | 95% | 0 | 94.46 | CP082199.1 |

Supplementary Table S7. Prevalence of ARGs in different genomic elements

| CARD ARG ID | NCBI Plasmid | NCBI WGS | NCBI Chromosome | NCBI Genomic Island |
|---|---|---|---|---|
| ARO:3002626 | 15.79 | 2.26 | 0 | 0 |
| ARO:3000410 | 10.97 | 7.21 | 1.52 | 0 |
| ARO:3002693 | 4.38 | 0.95 | 0.76 | 1.66 |
| ARO:3002660 | 3.9 | 7.53 | 4.55 | 33.33 |
| ARO:3000319 | 3.57 | 8.58 | 2.53 | 0 |
| ARO:3000251 | 3.31 | 8.62 | 2.62 | 0.86 |
| ARO:3003741 | 3.17 | 31.86 | 35.75 | 0 |
| ARO:3003109 | 3.12 | 31.73 | 35.93 | 4.4 |
| ARO:3002647 | 3.01 | 11.49 | 8.12 | 5.73 |
| ARO:3002897 | 2.86 | 9.81 | 4.54 | 6.02 |
| ARO:3005010 | 2.85 | 6.52 | 0 | 0 |
| ARO:3001418 | 2.7 | 16.9 | 45.31 | 0.63 |
| ARO:3000858 | 2.5 | 1.32 | 1.83 | 0 |
| ARO:3000191 | 1.64 | 0.2 | 0 | 0 |
| ARO:3002847 | 1.09 | 1.67 | 1.24 | 1.89 |
| ARO:3002695 | 0.94 | 1.51 | 1.24 | 1.89 |
| ARO:3005100 | 0.91 | 6.88 | 0 | 0 |
| ARO:3002865 | 0.67 | 3.32 | 1.4 | 1.43 |
| ARO:3003693 | 0.58 | 13.86 | 29.29 | 0 |
| ARO:3003710 | 0.58 | 38.88 | 58.59 | 0 |
| ARO:3003746 | 0.49 | 0 | 0 | 0 |
| ARO:3004621 | 0.42 | 4.82 | 1.77 | 0 |
| ARO:3002602 | 0.42 | 0.36 | 0.35 | 2.52 |
| ARO:3003200 | 0.36 | 3.92 | 2.48 | 0 |
| ARO:3003692 | 0.29 | 28.29 | 42.02 | 0 |
| ARO:3003699 | 0.29 | 65.99 | 97.7 | 0 |
| ARO:3001917 | 0.26 | 0.6 | 0.06 | 0 |
| ARO:3001889 | 0.16 | 0.01 | 0 | 0 |
| ARO:3000186 | 0.16 | 0.03 | 0.18 | 0.63 |
| ARO:3003013 | 0.16 | 0.24 | 0 | 0 |
| ARO:3002628 | 0.11 | 3.17 | 4.89 | 4.87 |
| ARO:3003107 | 0.11 | 0.1 | 0 | 0 |
| ARO:3000780 | 0.1 | 57.83 | 87.96 | 0 |
| ARO:3000782 | 0.1 | 61.75 | 93.81 | 0 |
| ARO:3000781 | 0.1 | 62.95 | 96.99 | 0 |
| ARO:3000559 | 0.1 | 42.32 | 71.86 | 0 |
| ARO:3000413 | 0.06 | 0.3 | 0 | 0 |
| ARO:3004573 | 0.05 | 40.25 | 64.42 | 0 |
| ARO:3000778 | 0.05 | 46.63 | 59.29 | 0 |
| ARO:3000620 | 0.05 | 50.45 | 79.82 | 0 |
| ARO:3000779 | 0.05 | 50.09 | 72.39 | 0 |
| ARO:3000753 | 0.05 | 2.98 | 5.49 | 0 |
| ARO:3004574 | 0.05 | 50.82 | 77.88 | 0 |
| ARO:3000774 | 0.05 | 40.59 | 50.62 | 0 |
| ARO:3000553 | 0.05 | 59.56 | 84.6 | 0.63 |

| | | | | |
|---|---|---|---|---|
| ARO:3000375 | 0.05 | 0.01 | 0.18 | 0 |
| ARO:3002972 | 0.04 | 56.09 | 99.74 | 0 |
| ARO:3000792 | 0.01 | 6.34 | 10.98 | 0 |
| ARO:3000904 | 0 | 0.02 | 0 | 0 |
| ARO:3000616 | 0 | 0.35 | 0.49 | 0 |
| ARO:3000522 | 0 | 1.05 | 3.57 | 0 |
| ARO:3004042 | 0 | 1.6 | 3.57 | 0 |
| ARO:3000216 | 0 | 73.41 | 85.05 | 0 |
| ARO:3002671 | 0 | 0 | 0 | 9.09 |
| ARO:3004039 | 0 | 1.18 | 5.91 | 0 |
| ARO:3003002 | 0 | 5.04 | 3.85 | 9.09 |
| ARO:3000190 | 0 | 0.04 | 0 | 0 |
| ARO:3002837 | 0 | 0.04 | 0 | 0 |
| ARO:3000196 | 0 | 0.02 | 0.49 | 0 |
| ARO:3002597 | 0 | 2.4 | 0 | 0 |
| ARO:3002909 | 0 | 0.17 | 0 | 0 |
| ARO:3003069 | 0 | 0 | 0.93 | 0 |
| ARO:3002965 | 0 | 0.04 | 0 | 0 |
| ARO:3002926 | 0 | 0 | 0.93 | 0 |
| ARO:3004253 | 0 | 11.11 | 0 | 0 |
| ARO:3002688 | 0 | 0.06 | 0 | 0 |
| ARO:3004442 | 0 | 0.05 | 0 | 0 |
| ARO:3004659 | 0 | 15.12 | 26.92 | 0 |
| ARO:3000777 | 0 | 55.81 | 100 | 50 |
| ARO:3000768 | 0 | 82.61 | 100 | 0 |
| ARO:3005051 | 0 | 0.53 | 2.12 | 0 |
| ARO:3001613 | 0 | 2.91 | 3.72 | 0 |
| ARO:3000775 | 0 | 0 | 0 | 0.63 |
| ARO:3000549 | 0 | 0.2 | 0.35 | 0 |
| ARO:3006346 | 0 | 0.2 | 0.18 | 0 |
| ARO:3004577 | 0 | 3.99 | 6.73 | 0 |
| ARO:3002639 | 0 | 0.97 | 0 | 0 |
| ARO:3000412 | 0 | 7.14 | 0 | 0 |
| ARO:3002369 | 0 | 1.33 | 0 | 0 |
| ARO:3000621 | 0 | 0.01 | 0 | 0 |
| ARO:3000194 | 0 | 80 | 100 | 100 |
| ARO:3001060 | 0 | 0.02 | 0 | 0 |
| ARO:3002683 | 0 | 4.34 | 0 | 0 |
| ARO:3000935 | 0 | 0 | 0 | 0.63 |
| ARO:3001071 | 0 | 0.14 | 0.18 | 0 |
| ARO:3000316 | 0 | 0.01 | 0 | 0 |
| ARO:3002859 | 0 | 0 | 0.18 | 0 |
| ARO:3004089 | 0 | 0 | 100 | 0 |
| ARO:3005098 | 0 | 2.08 | 66.67 | 0 |
| ARO:3003839 | 0 | 1.6 | 0 | 0 |
| ARO:3002868 | 0 | 43.59 | 36.67 | 6.67 |
| ARO:3002838 | 0 | 6.25 | 0 | 0 |

| | | | | |
|---|---|---|---|---|
| ARO:3000567 | 0 | 0 | 0 | 8.33 |
| ARO:3007122 | 0 | 2.04 | 0 | 0 |
| ARO:3000979 | 0 | 0.13 | 0 | 0 |
| ARO:3005260 | 0 | 0.09 | 0 | 0 |
| ARO:3001878 | 0 | 0.16 | 0.35 | 0 |
| ARO:3001328 | 0 | 69.99 | 87.19 | 0 |
| ARO:3004056 | 0 | 22.68 | 0.61 | 0 |
| ARO:3002985 | 0 | 6.05 | 15.03 | 0 |
| ARO:3003698 | 0 | 19.93 | 39.42 | 0 |
| ARO:3003700 | 0 | 3.76 | 13.34 | 0 |
